## Supplementary material for "Dissecting surveying behavior of reactive microglia under chronic neurodegeneration": Figure S1

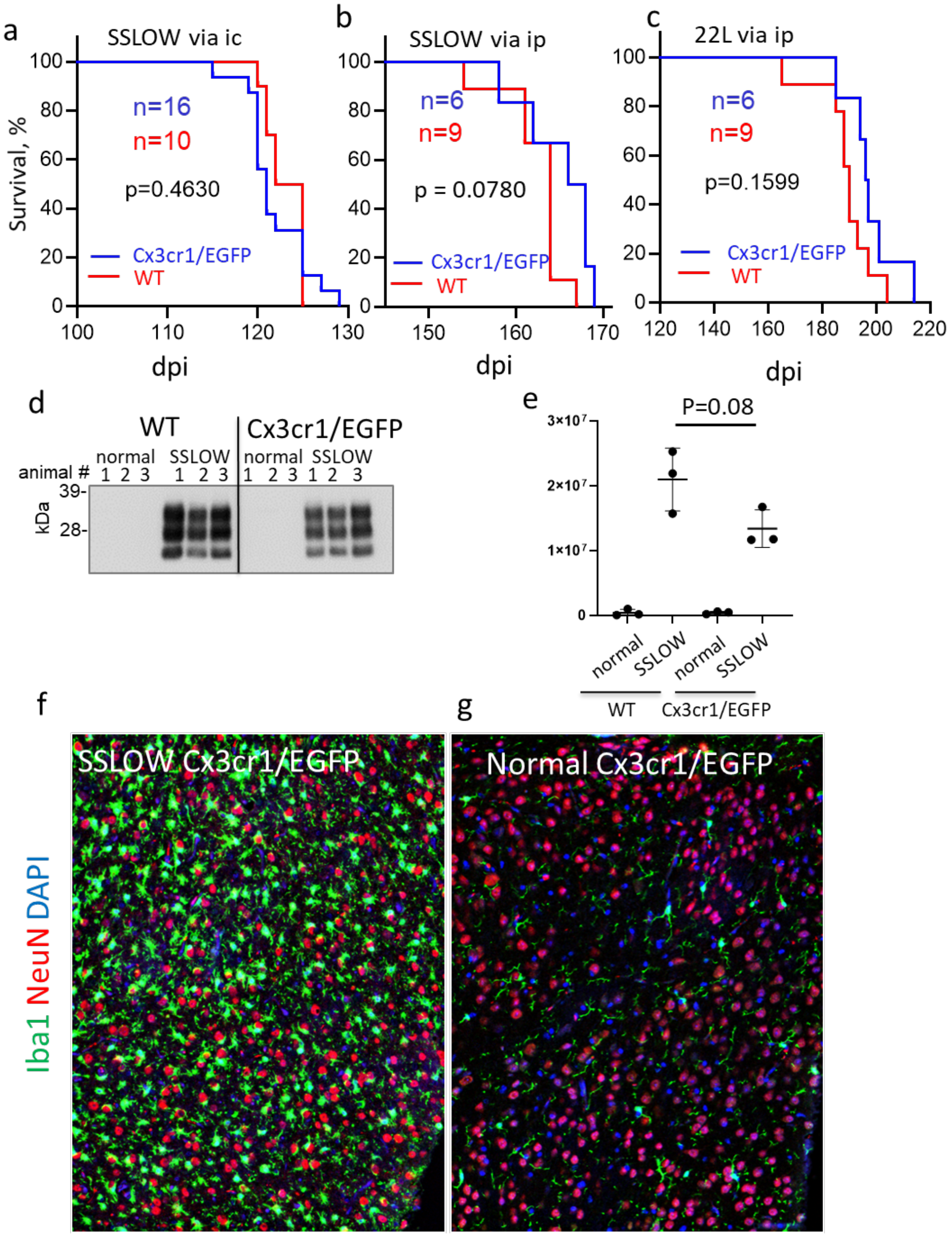


**Figure S1. Prion pathogenesis in not changed in Cx3cr1/EGFP mice. a,b,c** Survival curves for Cx3cr1/EGFP and WT (C57Bl/6J) mice inoculated with SSLOW via ic route (**a**), SSLOW via ip route (**b**) or 22L via ip route (**c**). Comparison by Mantel-Cox test. **d,e** Representative Western blot image (**d**) and quantification of $PrP^{Sc}$ (**e**) in brains of WT and Cx3cr1/EGFP mice infected via ip route. The data presented as Means ± SD; p by Brown-Forsythe and Welch ANOVA with Dunnett's multiple comparison test, n=3 per group. Data for non-infected WT and Cx3cr1/EGFP brains (normal) are shown as a reference. **f, g** Immunostaining for microglia (Iba1, green) and neurons (NeuN, red) showing reactive Iba1+ cells enveloping neurons in cortex of Cx3cr1/EGFP mice infected by SSLOW via ip route at the terminal stage of the disease (**f**); and lack of neuronal envelopment in adult, non-infected Cx3cr1/EGFP mice (**g**).

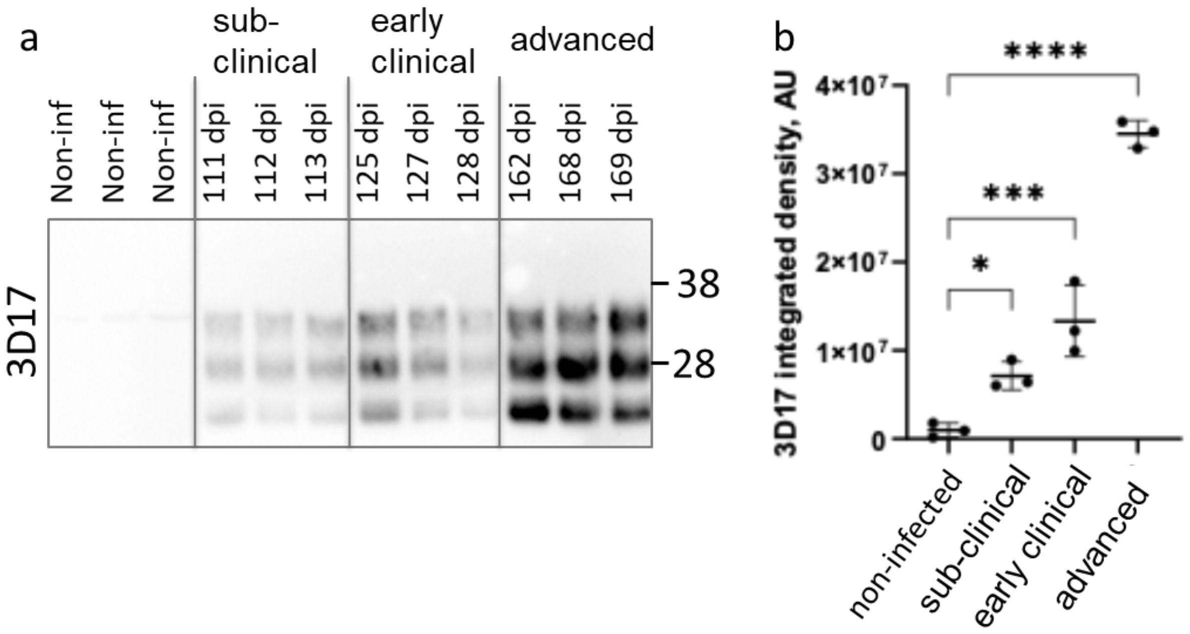


**Figure S2. Accumulation of PrP$^{Sc}$ in SSLOW-infected Cx3cr1/EGFP mice. a,** Western blot analysis of PrP$^{Sc}$ in brain homogenates from Cx3cr1/EGFP mice used for time-lapse imaging experiments, compared to non-infected controls. Mice were infected with SSLOW via intraperitoneal injection and analyzed at subclinical (111–113 dpi), early clinical (125–128 dpi), and advanced (162–169 dpi) disease stages. Non-infected mice were analyzed at 160–164 days of age. Samples were digested with Proteinase K and probed with the 3D17 anti-PrP antibody. **b**, Densitometric quantification of PrP$^{Sc}$ signal intensity. Data are shown as mean ± SD. Statistical significance was determined by ordinary one-way ANOVA with Dunnett's multiple comparison test (*$p < 0.05$, ***$p < 0.001$, ****$p < 0.0001$). *N* = 3 mice per group.

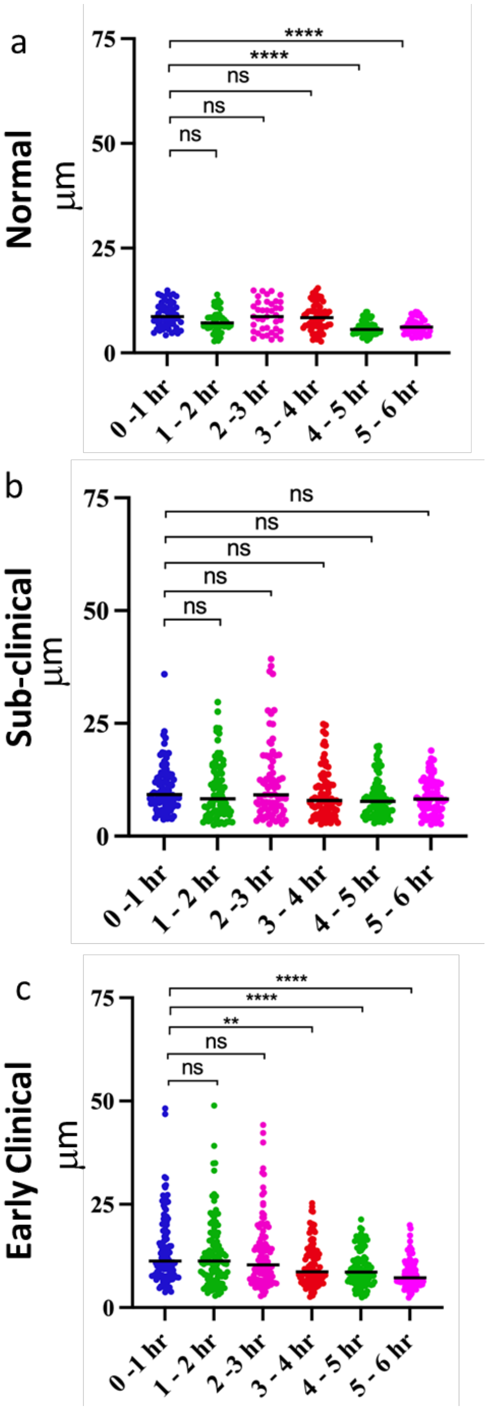


**Figure S3. Mobility of EGFP+ cells across six consecutive one-hour intervals post-slicing.** Acute cerebral cortical slices were prepared acutely using non-infected Cx3cr1/EGFP (normal) mice or Cx3cr1/EGFP mice infected with SSLOW via ip route at sub-clinical and early clinical stages of the disease. Distance covered by individual EGFP+ cells in one-hour periods across six consecutive time intervals. Means are marked by black lines. N=3 animals per group; n=40-65 cells per group, **$p<0.01$, ****$p<0.0001$, ns - non-significant by non-parametric Kruskal-Wallis test with Dunn's multiple comparison test.

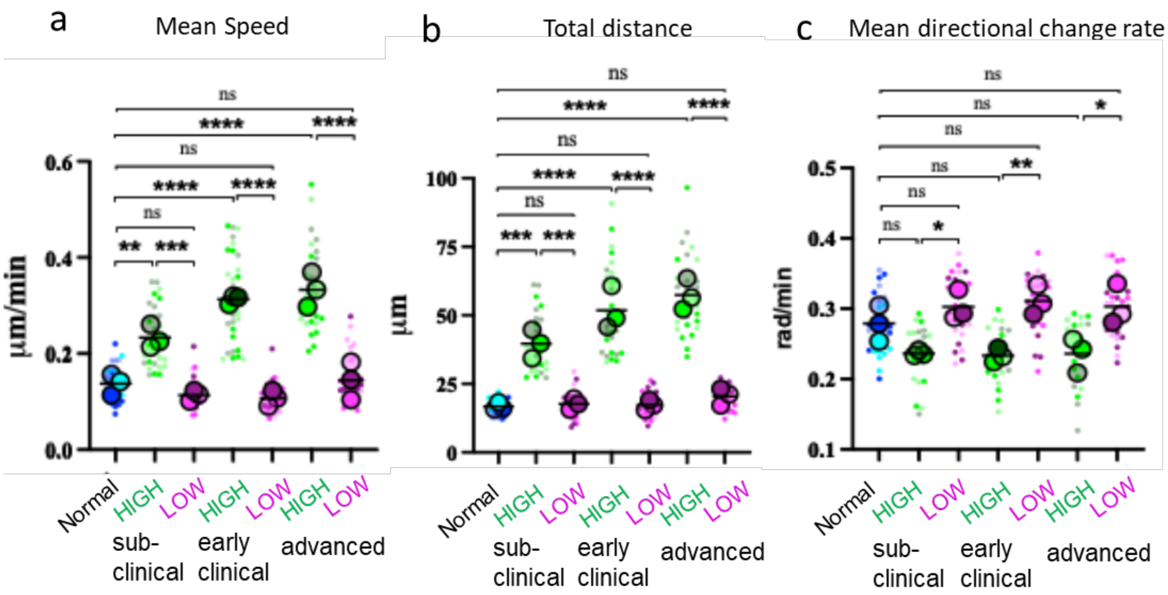


**Figure S4. Inter-animal variation in high- and low-mobility behavioral phenotypes.** Mean speed (**a**), total distance travelled over a 3-hour period (**b**), and mean directional change rate (**c**) were quantified for high- and low-mobility EGFP$^{+}$ cells in cerebral cortical slices from SSLOW-infected mice at three disease stages, and from non-infected control mice. In Superplots, colors indicate individual animals, dots represent single cells, circles show average value for each animal, black lines indicate group means calculated from biological replicates (animals). N=3 animals per group. Significance was determined by ordinary one-way ANOVA: *p<0.05, **p<0.01, ***p<0.001, ****p<0.0001, ns - non-significant.

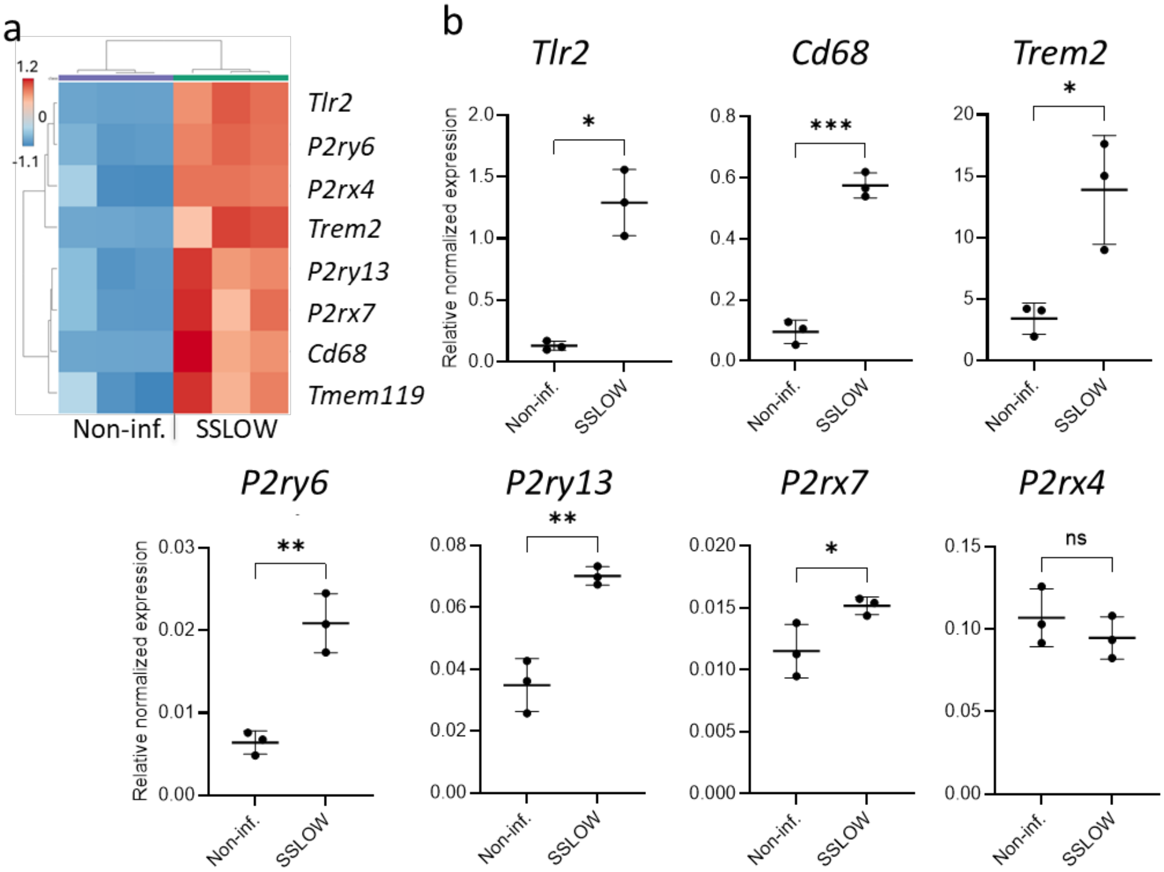


**Figure S5. Gene expression analysis in SSLOW-infected mice. a,** Heatmap depicting differential gene expression in bulk brain tissue from SSLOW-infected C57BL/6J mice at advanced disease stage. $N = 3$ mice. **b**, Relative gene expression in SSLOW-infected versus age-matched non-infected C57BL/6J mice, normalized by TMEM119 expression. Data are presented as mean ± SD. Statistical significance was assessed using unpaired *t*-test with Welch's correction (*$p < 0.05$, **$p < 0.01$, ***$p < 0.001$). $N = 3$ mice per group.

.

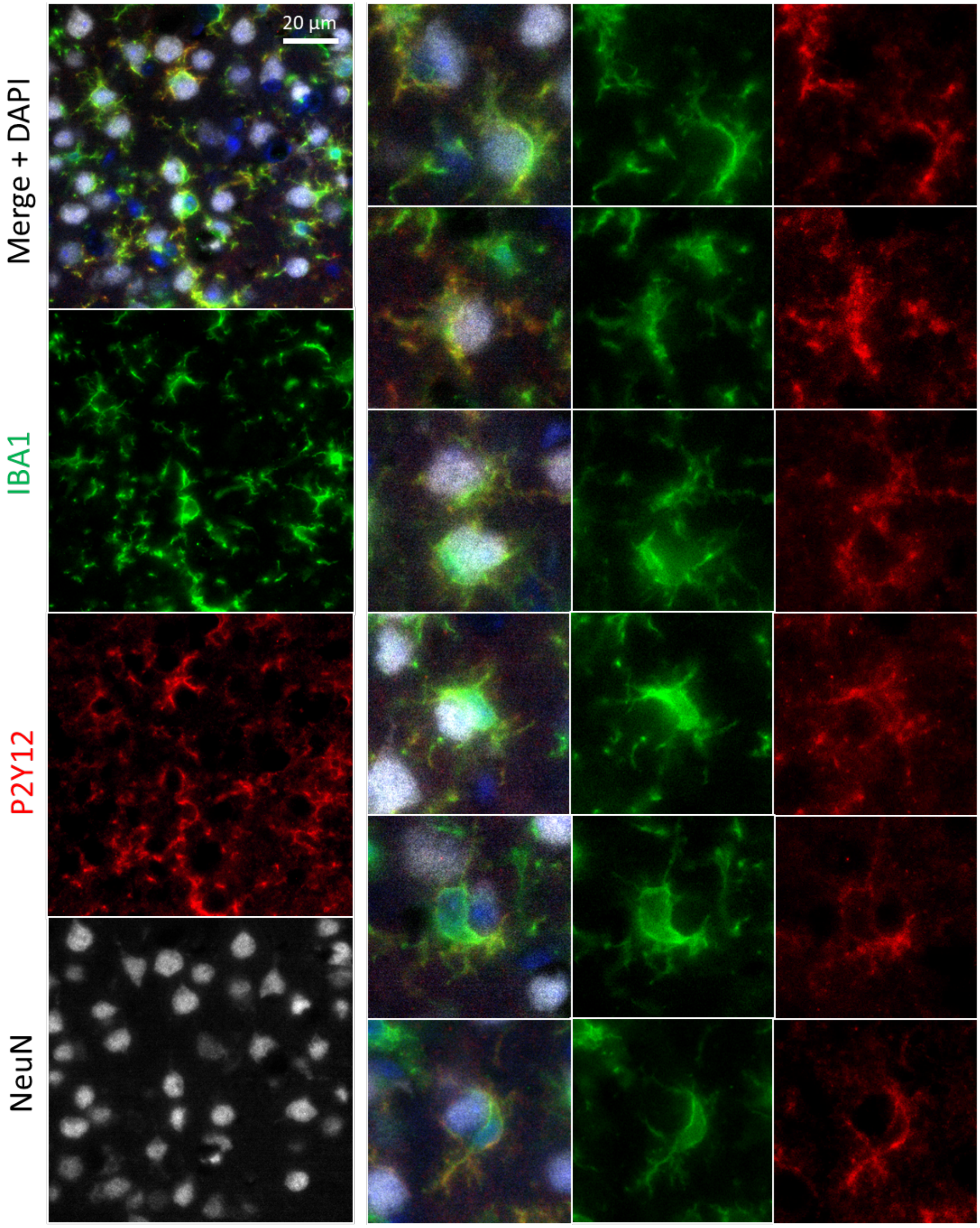


**Figure S6. Envelopment of neurons by P2Y12-positive microglia.** Representative images showing envelopment of neurons by reactive microglia in the cerebral cortex of SSLOW-infected Cx3cr1/EGFP mice (ip inoculation). Brain sections were immunostained with anti-IBA1 (green), anti-P2Y12 (red), and anti-NeuN (gray) antibodies. Right panels present a gallery of envelopment events.

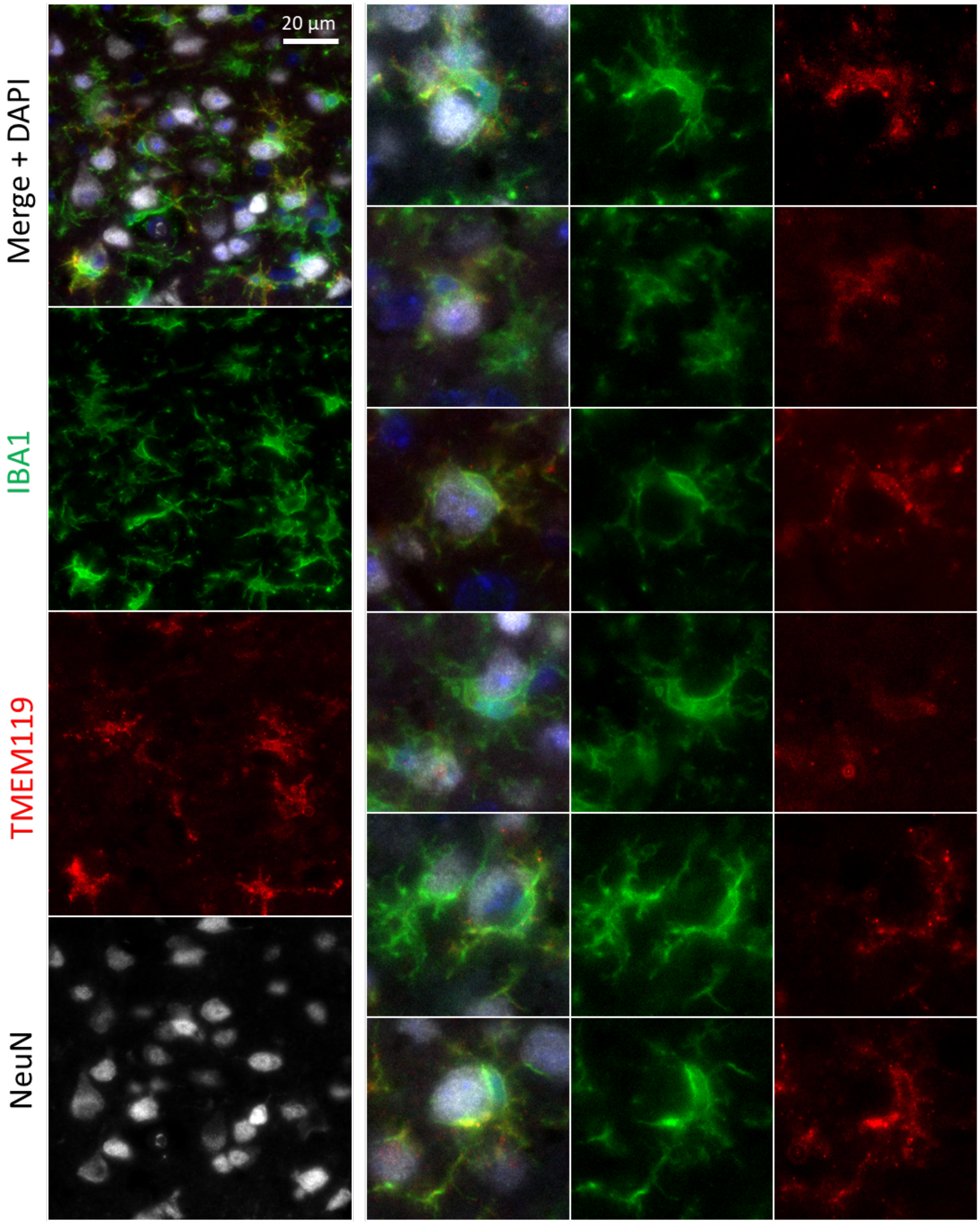


**Figure S7. Envelopment of neurons by TMEM119-positive microglia.** Representative images showing envelopment of neurons by reactive microglia in the cerebral cortex of SSLOW-infected Cx3cr1/EGFP mice (ip inoculation). Brain sections were immunostained with anti-IBA1 (green), anti-TMEM119 (red), and anti-NeuN (gray) antibodies. Right panels present a gallery of envelopment events.
