## Supplementary material for "Dissecting surveying behavior of reactive microglia under chronic neurodegeneration": Video S1

### Slide 1
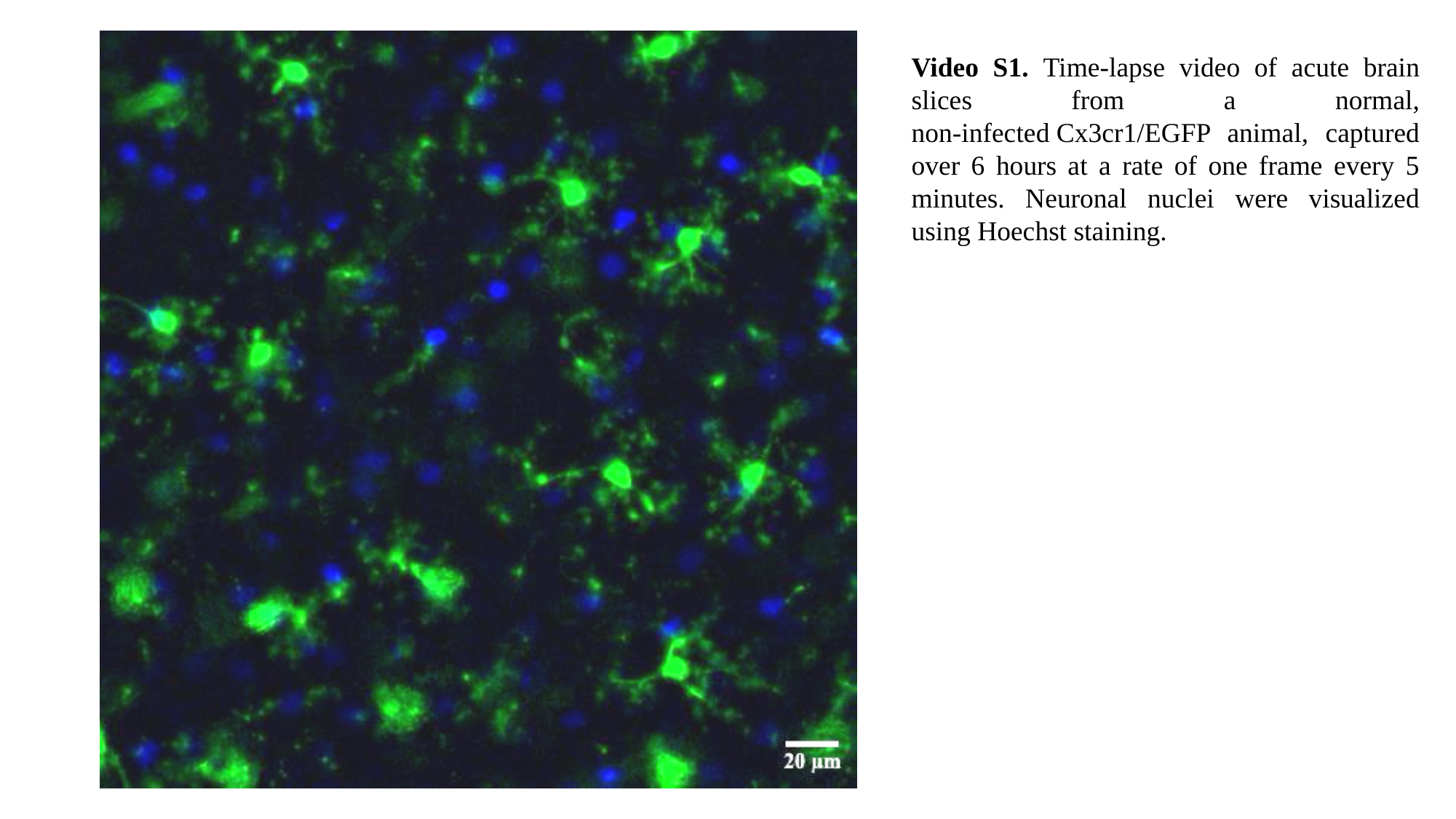

Video S1. Time-lapse video of acute brain slices from a normal, non-infected Cx3cr1/EGFP animal, captured over 6 hours at a rate of one frame every 5 minutes. Neuronal nuclei were visualized using Hoechst staining.

### Slide 2
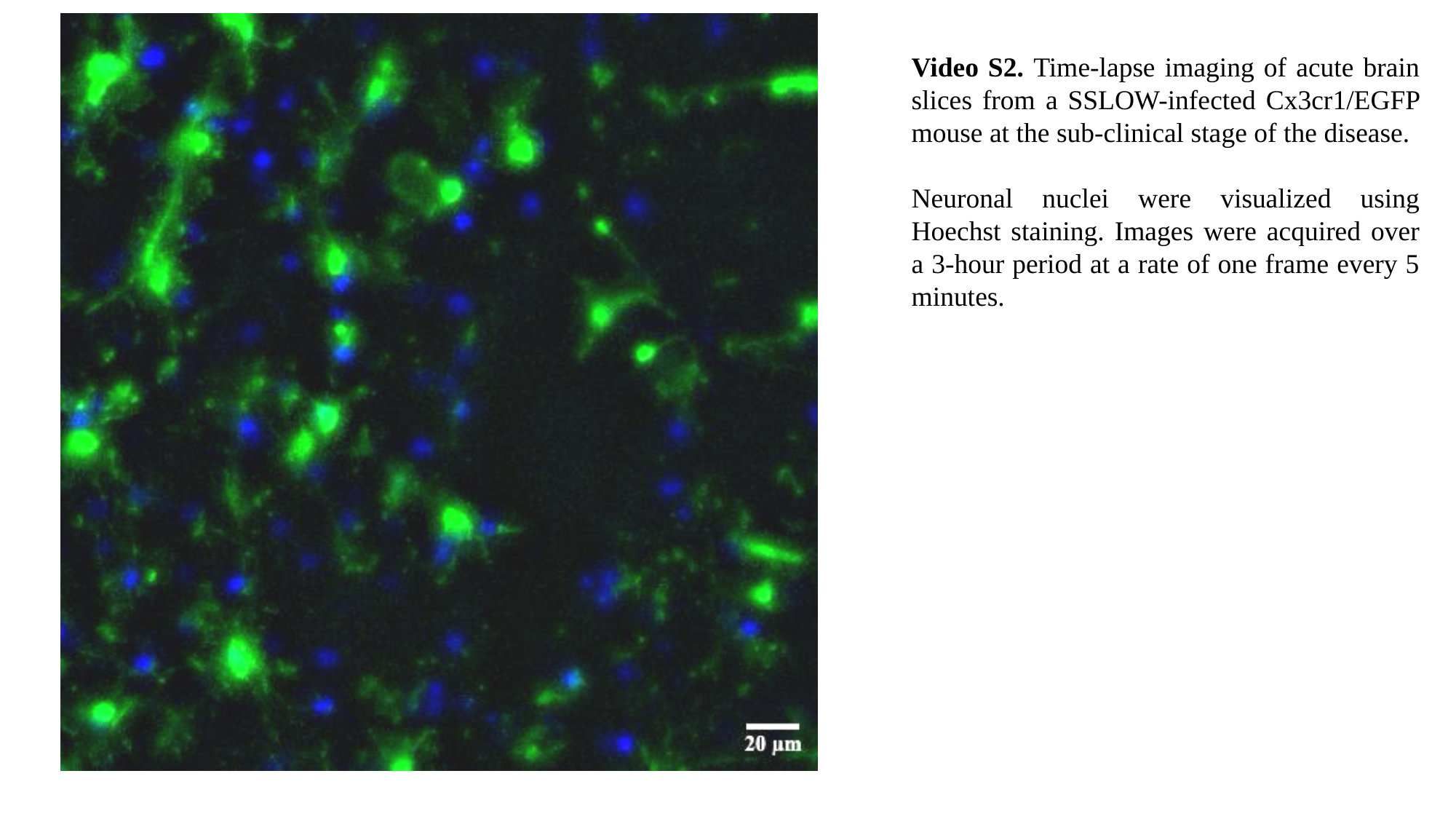

Video S2. Time-lapse imaging of acute brain slices from a SSLOW-infected Cx3cr1/EGFP mouse at the sub-clinical stage of the disease.
Neuronal nuclei were visualized using Hoechst staining. Images were acquired over a 3-hour period at a rate of one frame every 5 minutes.

### Slide 3
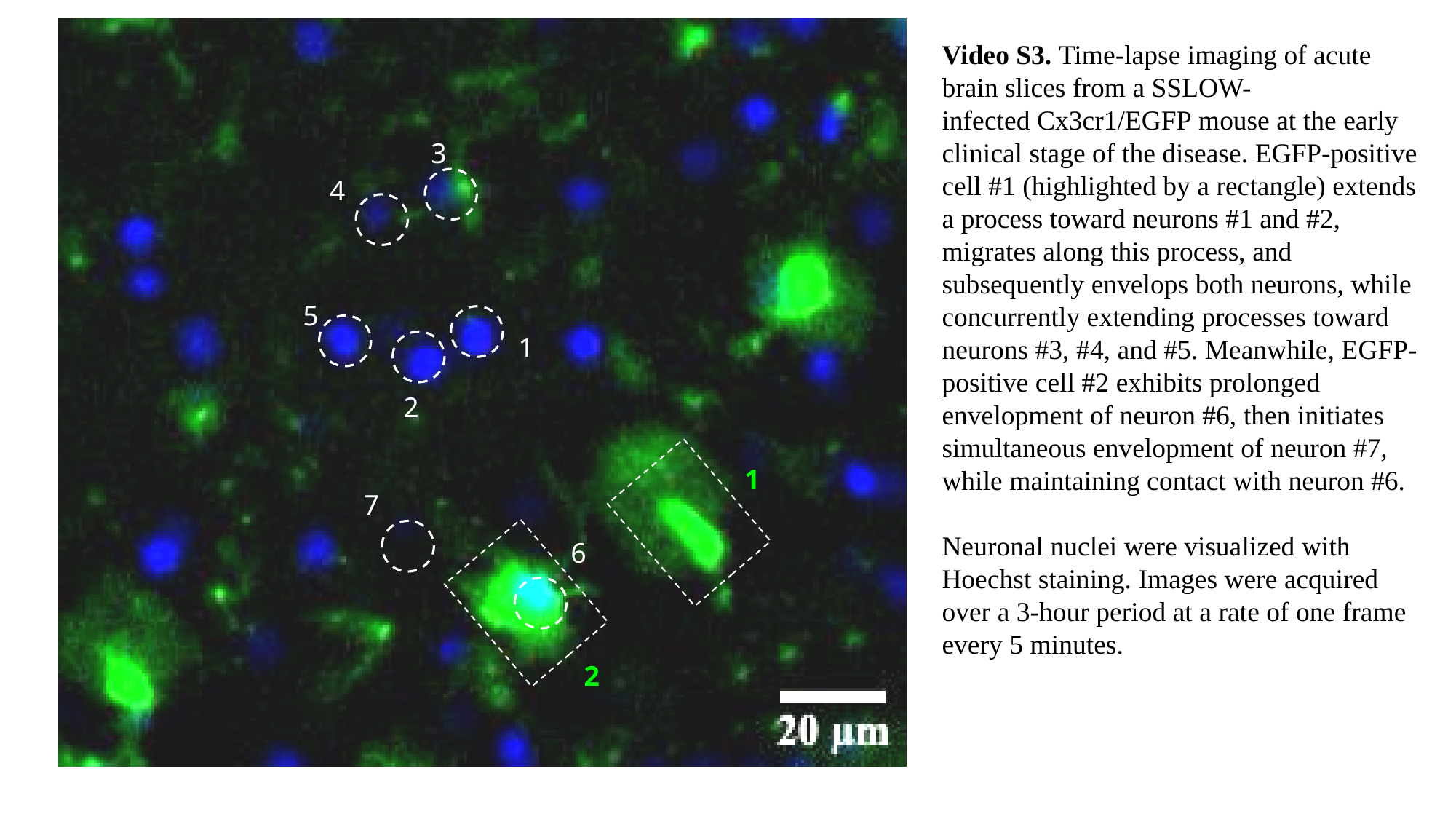

3
1
2
1
7
6
2
Video S3. Time-lapse imaging of acute brain slices from a SSLOW-infected Cx3cr1/EGFP mouse at the early clinical stage of the disease. EGFP-positive cell #1 (highlighted by a rectangle) extends a process toward neurons #1 and #2, migrates along this process, and subsequently envelops both neurons, while concurrently extending processes toward neurons #3, #4, and #5. Meanwhile, EGFP-positive cell #2 exhibits prolonged envelopment of neuron #6, then initiates simultaneous envelopment of neuron #7, while maintaining contact with neuron #6.
Neuronal nuclei were visualized with Hoechst staining. Images were acquired over a 3-hour period at a rate of one frame every 5 minutes.
4
5

### Slide 4
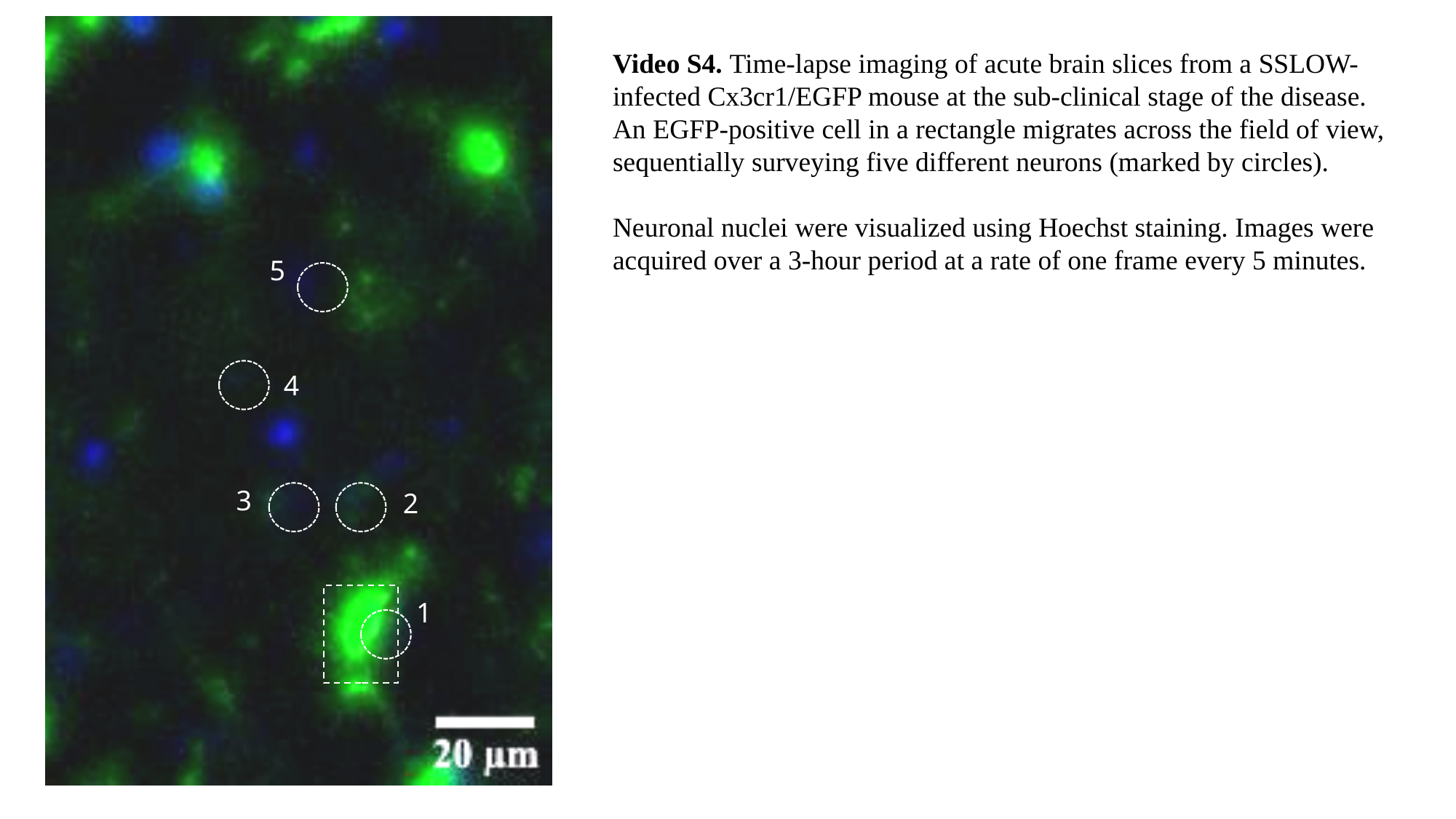

5
4
3
2
1
Video S4. Time-lapse imaging of acute brain slices from a SSLOW-infected Cx3cr1/EGFP mouse at the sub-clinical stage of the disease. An EGFP-positive cell in a rectangle migrates across the field of view, sequentially surveying five different neurons (marked by circles).
Neuronal nuclei were visualized using Hoechst staining. Images were acquired over a 3-hour period at a rate of one frame every 5 minutes.

### Slide 5
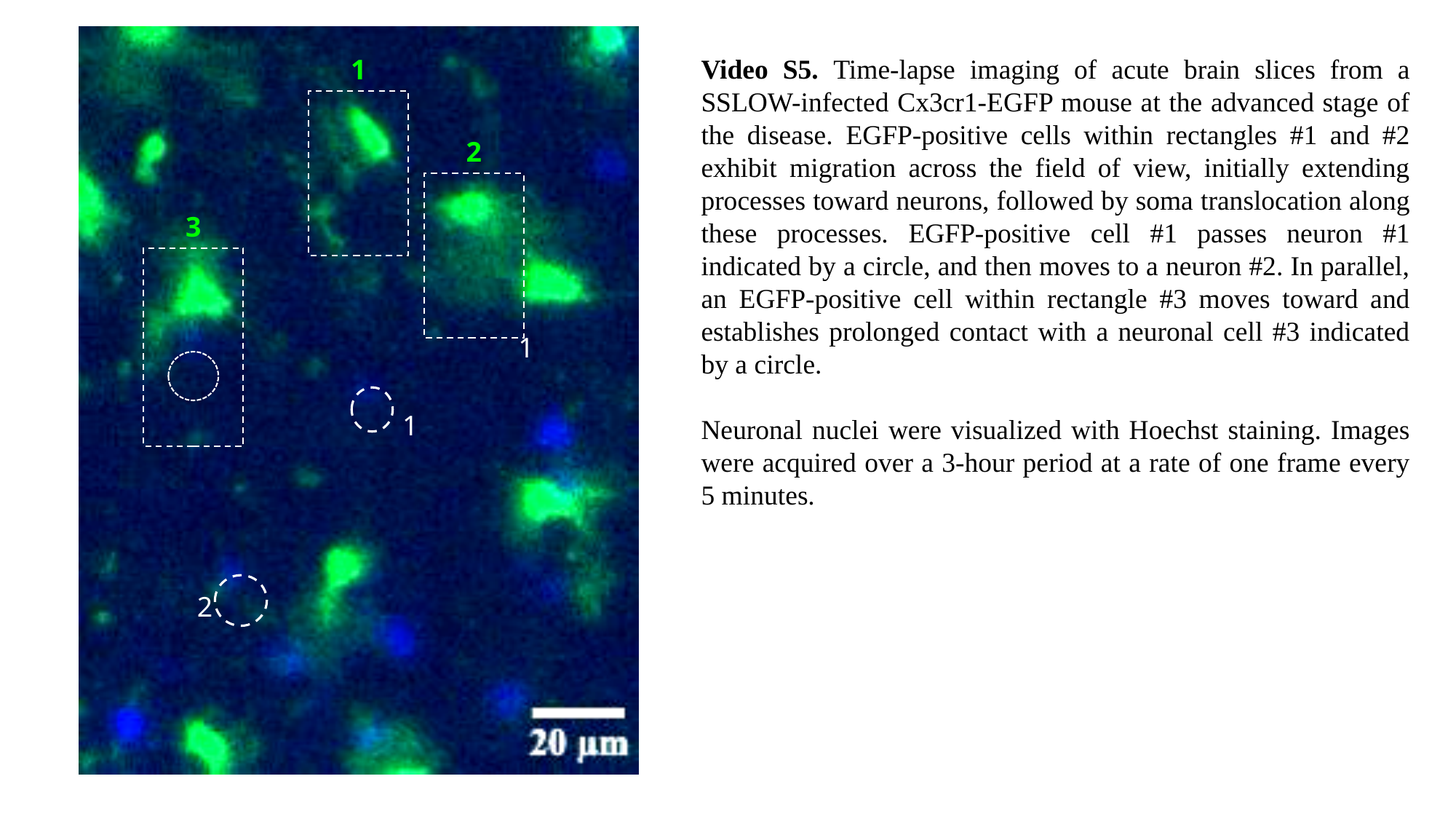

1
2
3
Video S5. Time-lapse imaging of acute brain slices from a SSLOW-infected Cx3cr1-EGFP mouse at the advanced stage of the disease. EGFP-positive cells within rectangles #1 and #2 exhibit migration across the field of view, initially extending processes toward neurons, followed by soma translocation along these processes. EGFP-positive cell #1 passes neuron #1 indicated by a circle, and then moves to a neuron #2. In parallel, an EGFP-positive cell within rectangle #3 moves toward and establishes prolonged contact with a neuronal cell #3 indicated by a circle.
Neuronal nuclei were visualized with Hoechst staining. Images were acquired over a 3-hour period at a rate of one frame every 5 minutes.
1
1
2

### Slide 6
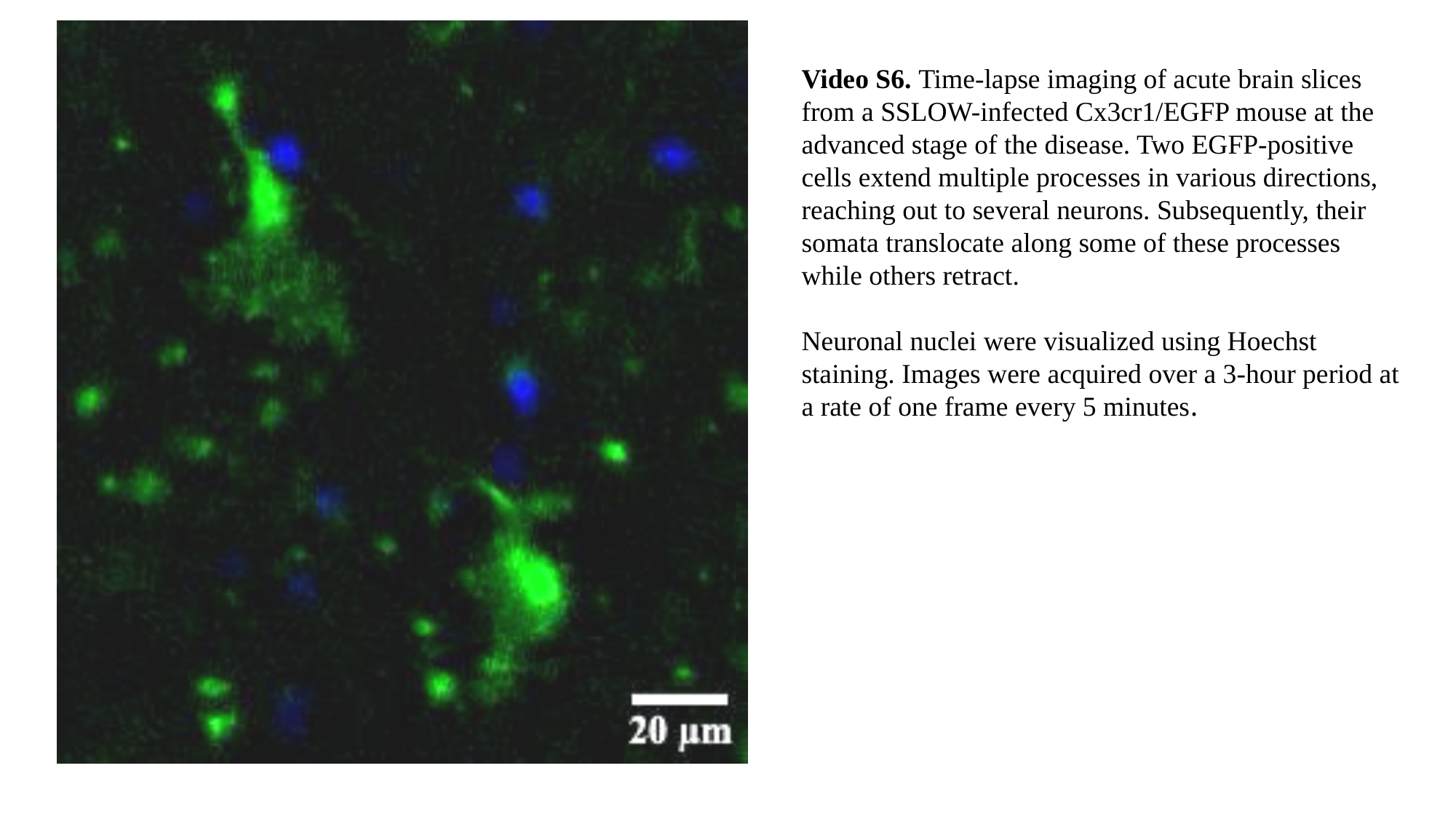

Video S6. Time-lapse imaging of acute brain slices from a SSLOW-infected Cx3cr1/EGFP mouse at the advanced stage of the disease. Two EGFP-positive cells extend multiple processes in various directions, reaching out to several neurons. Subsequently, their somata translocate along some of these processes while others retract.
Neuronal nuclei were visualized using Hoechst staining. Images were acquired over a 3-hour period at a rate of one frame every 5 minutes.

### Slide 7
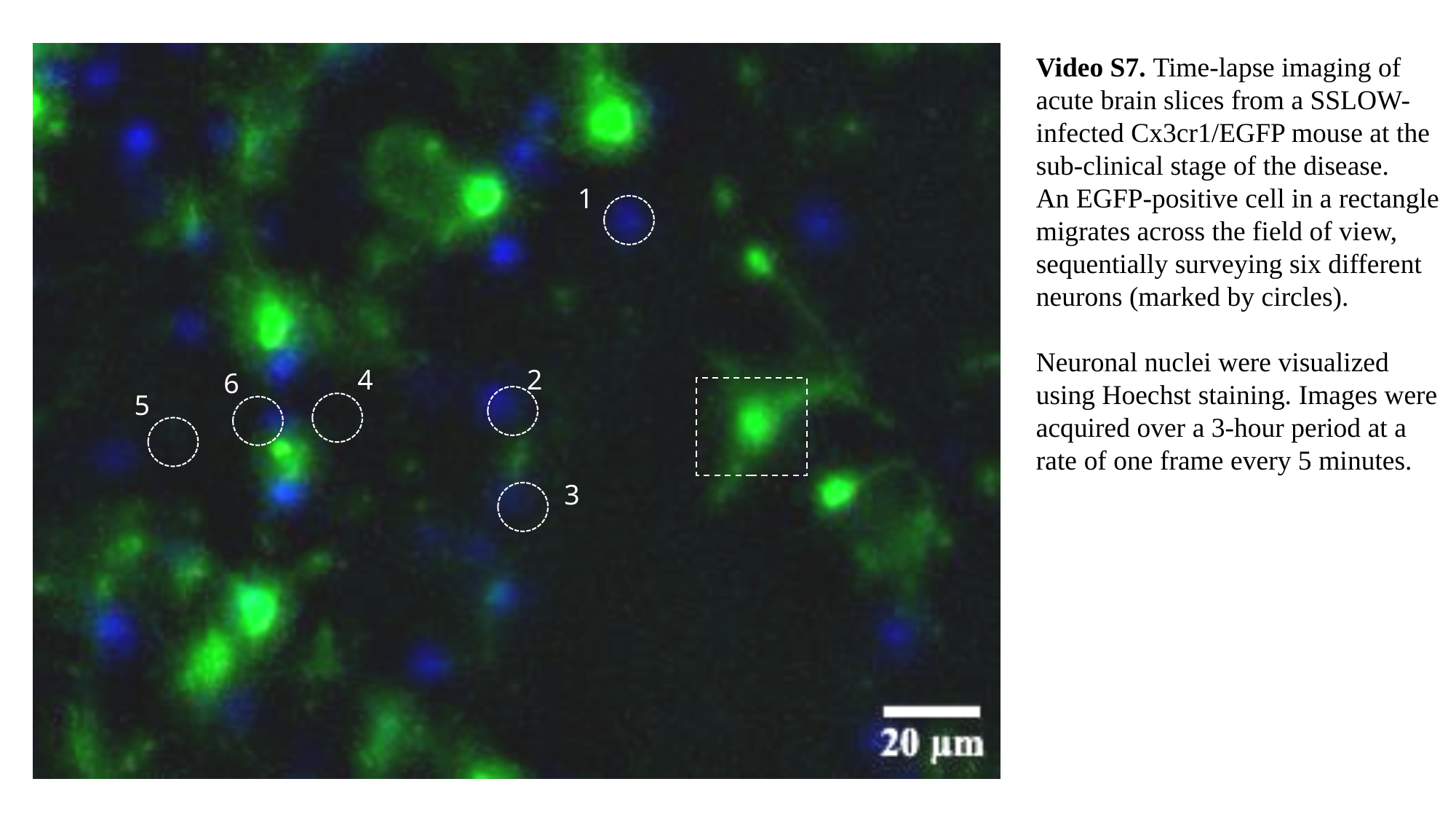

Video S7. Time-lapse imaging of acute brain slices from a SSLOW-infected Cx3cr1/EGFP mouse at the sub-clinical stage of the disease.
An EGFP-positive cell in a rectangle migrates across the field of view, sequentially surveying six different neurons (marked by circles).
Neuronal nuclei were visualized using Hoechst staining. Images were acquired over a 3-hour period at a rate of one frame every 5 minutes.
1
4
2
6
5
3

### Slide 8
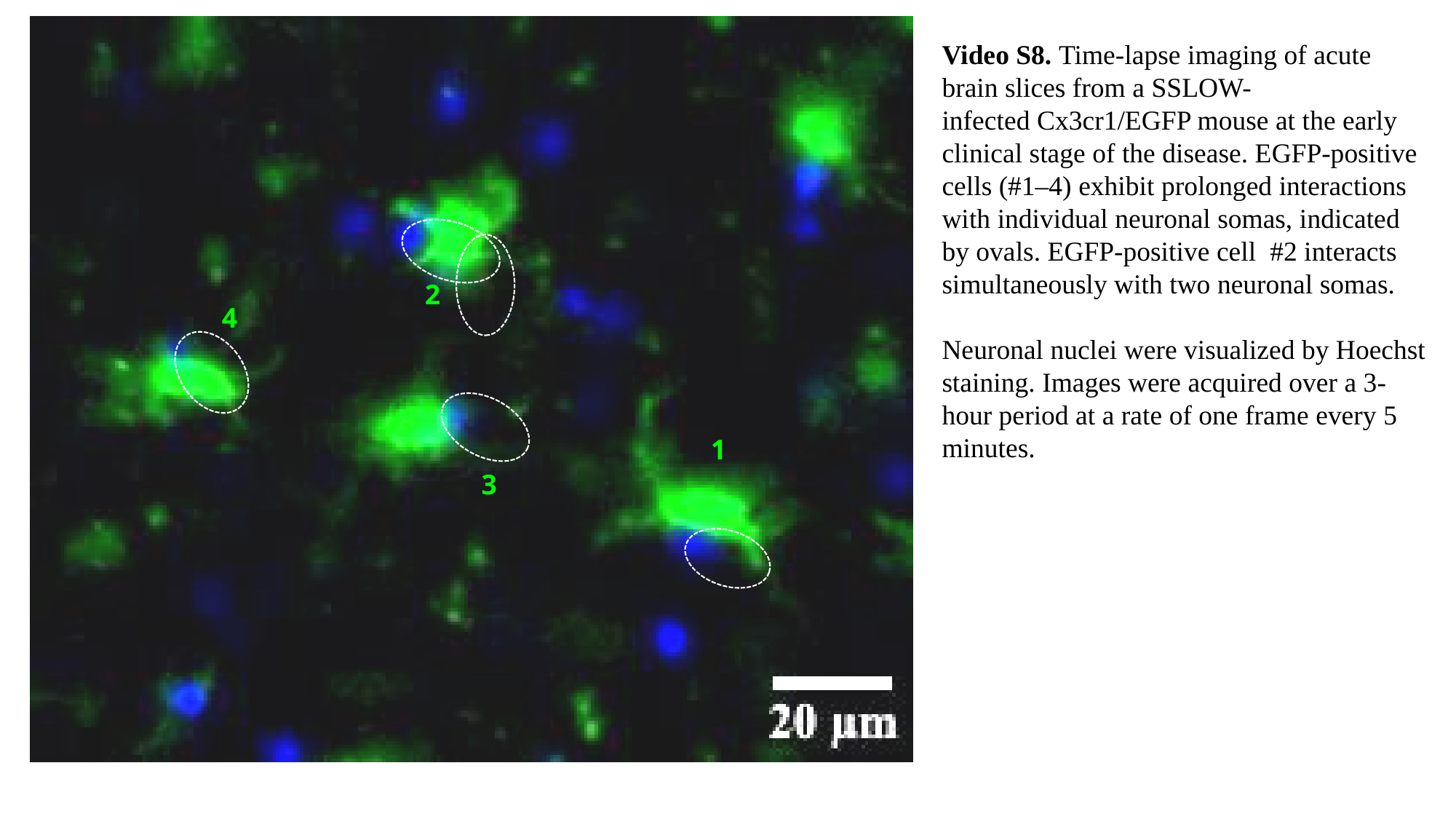

2
4
1
3
Video S8. Time-lapse imaging of acute brain slices from a SSLOW-infected Cx3cr1/EGFP mouse at the early clinical stage of the disease. EGFP-positive cells (#1–4) exhibit prolonged interactions with individual neuronal somas, indicated by ovals. EGFP-positive cell #2 interacts simultaneously with two neuronal somas.
Neuronal nuclei were visualized by Hoechst staining. Images were acquired over a 3-hour period at a rate of one frame every 5 minutes.

### Slide 9
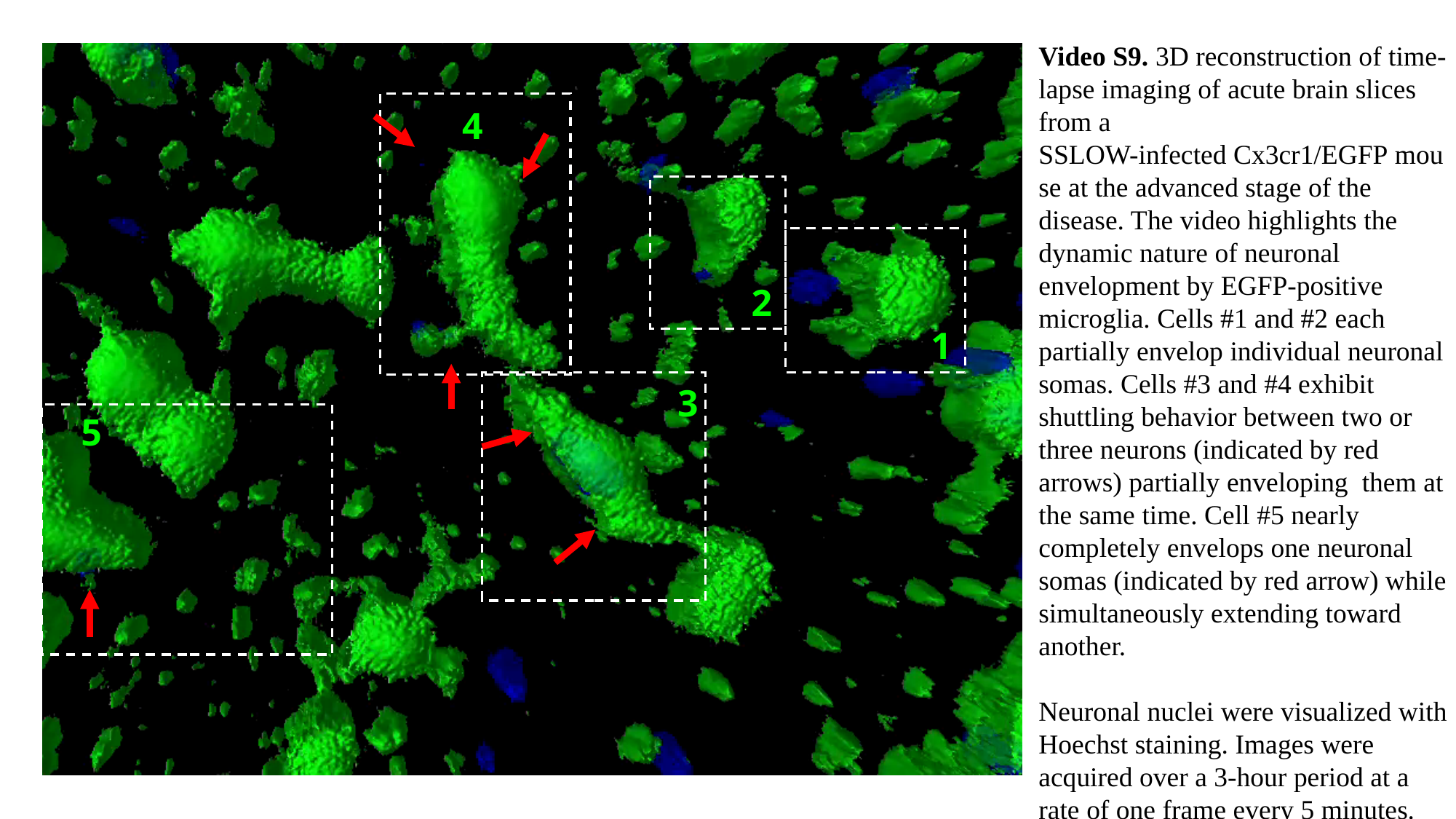

Video S9. 3D reconstruction of time-lapse imaging of acute brain slices from a SSLOW-infected Cx3cr1/EGFP mouse at the advanced stage of the disease. The video highlights the dynamic nature of neuronal envelopment by EGFP-positive microglia. Cells #1 and #2 each partially envelop individual neuronal somas. Cells #3 and #4 exhibit shuttling behavior between two or three neurons (indicated by red arrows) partially enveloping them at the same time. Cell #5 nearly completely envelops one neuronal somas (indicated by red arrow) while simultaneously extending toward another.
Neuronal nuclei were visualized with Hoechst staining. Images were acquired over a 3-hour period at a rate of one frame every 5 minutes.
4
2
1
3
5

### Slide 10
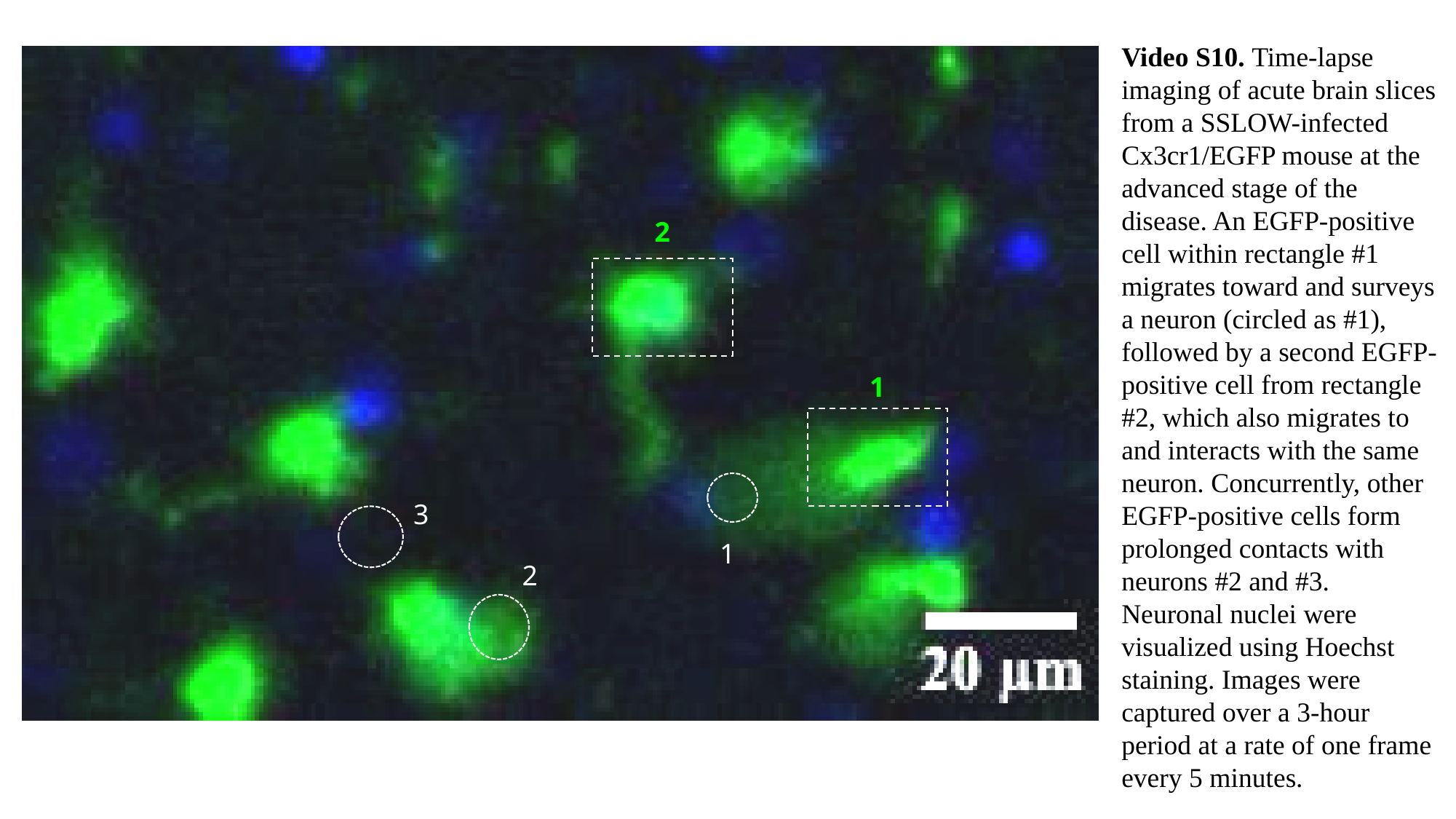

Video S10. Time-lapse imaging of acute brain slices from a SSLOW-infected Cx3cr1/EGFP mouse at the advanced stage of the disease. An EGFP-positive cell within rectangle #1 migrates toward and surveys a neuron (circled as #1), followed by a second EGFP-positive cell from rectangle #2, which also migrates to and interacts with the same neuron. Concurrently, other EGFP-positive cells form prolonged contacts with neurons #2 and #3.
Neuronal nuclei were visualized using Hoechst staining. Images were captured over a 3-hour period at a rate of one frame every 5 minutes.
2
1
3
1
2

### Slide 11
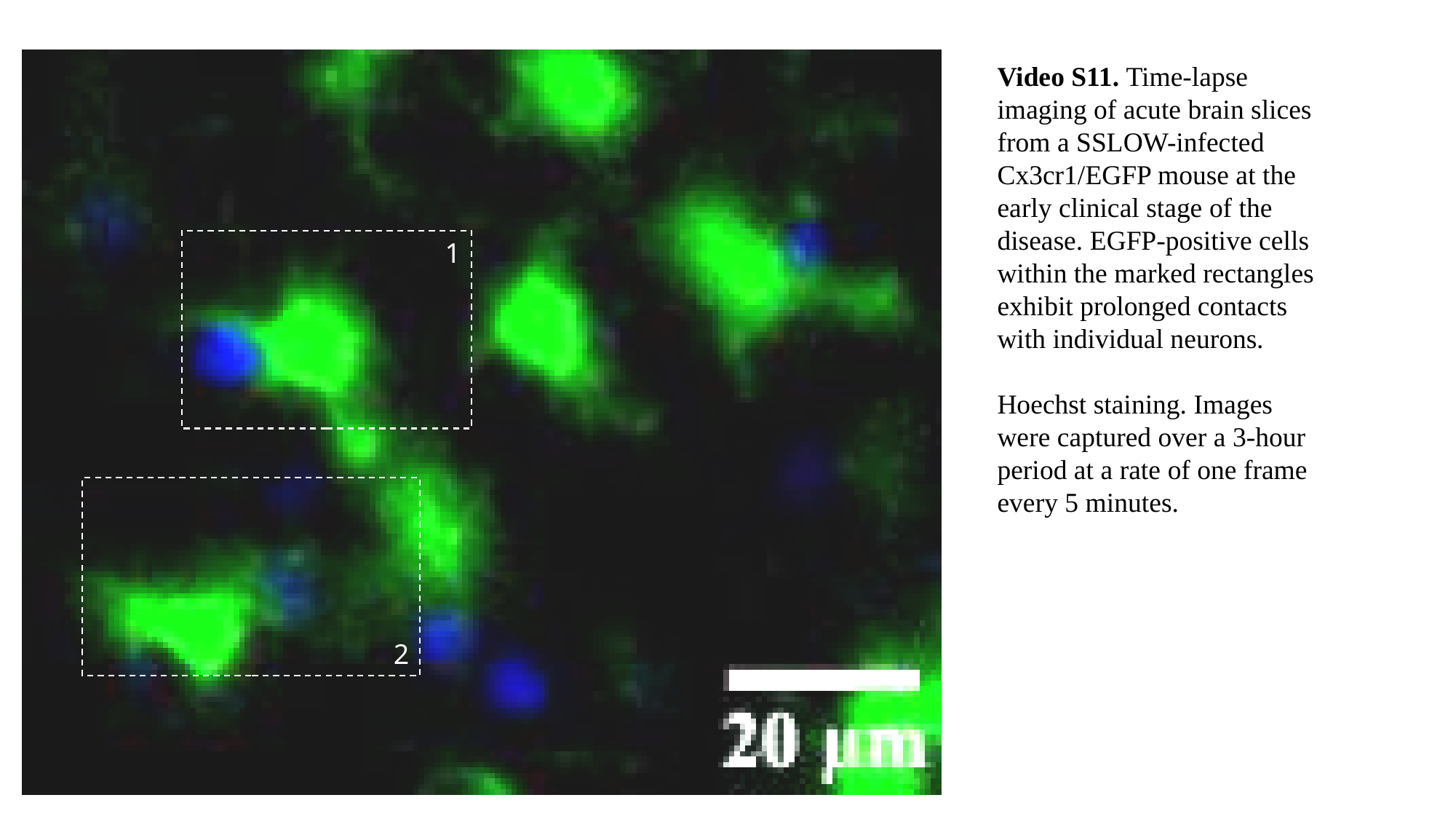

1
2
Video S11. Time-lapse imaging of acute brain slices from a SSLOW-infected Cx3cr1/EGFP mouse at the early clinical stage of the disease. EGFP-positive cells within the marked rectangles exhibit prolonged contacts with individual neurons.
Hoechst staining. Images were captured over a 3-hour period at a rate of one frame every 5 minutes.

### Slide 12
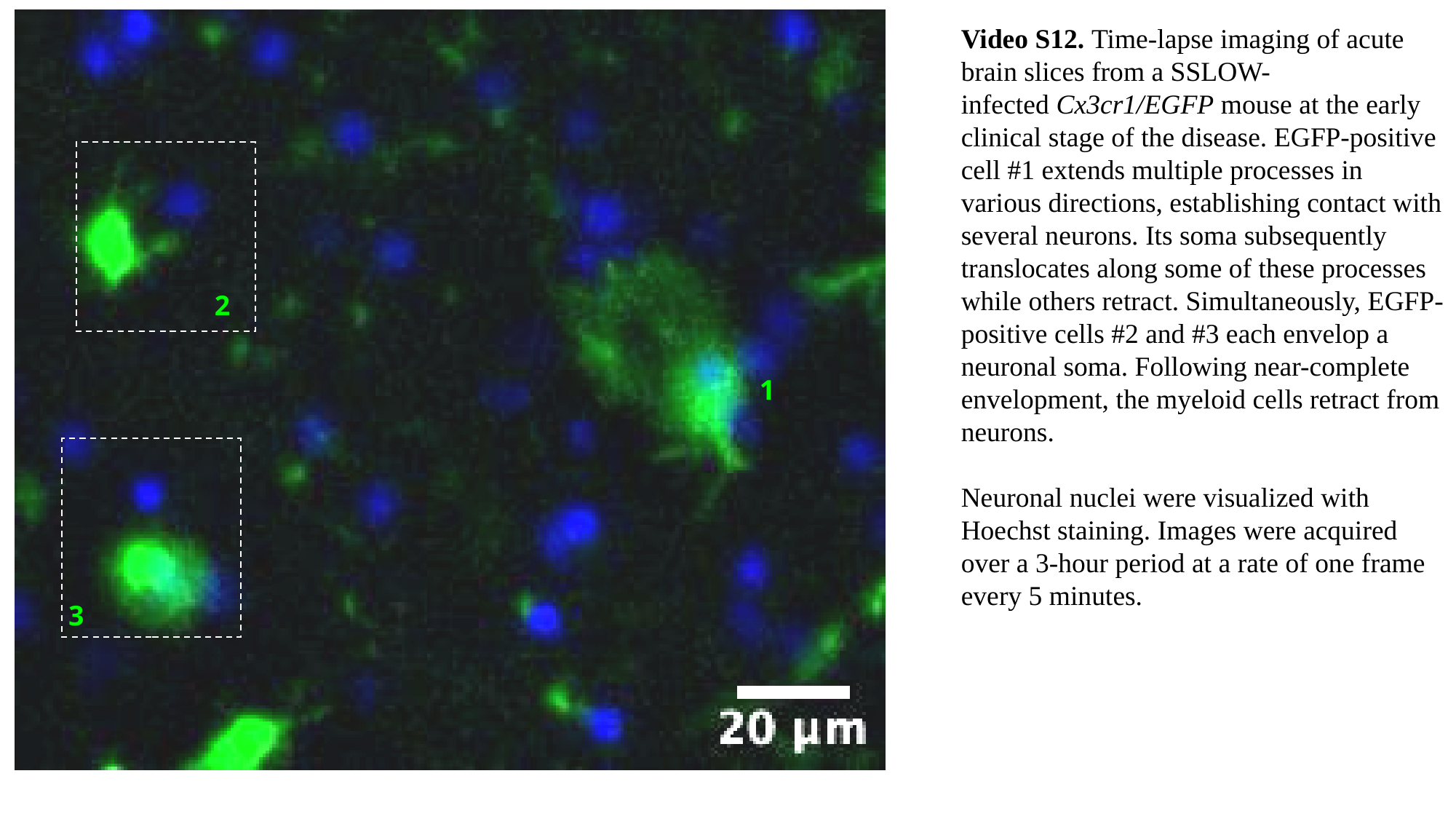

Video S12. Time-lapse imaging of acute brain slices from a SSLOW-infected Cx3cr1/EGFP mouse at the early clinical stage of the disease. EGFP-positive cell #1 extends multiple processes in various directions, establishing contact with several neurons. Its soma subsequently translocates along some of these processes while others retract. Simultaneously, EGFP-positive cells #2 and #3 each envelop a neuronal soma. Following near-complete envelopment, the myeloid cells retract from neurons.
Neuronal nuclei were visualized with Hoechst staining. Images were acquired over a 3-hour period at a rate of one frame every 5 minutes.
2
1
3

### Slide 13
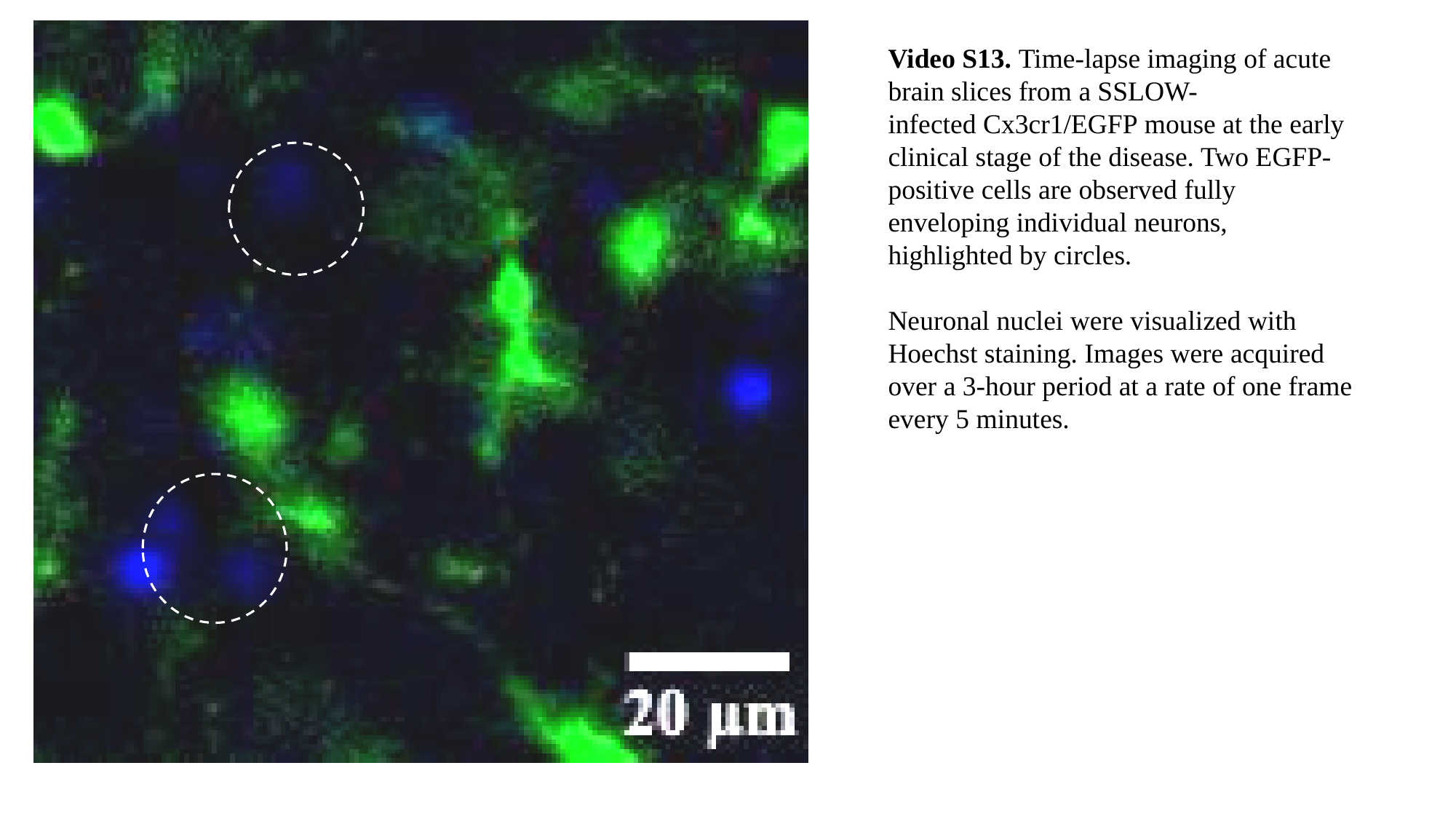

Video S13. Time-lapse imaging of acute brain slices from a SSLOW-infected Cx3cr1/EGFP mouse at the early clinical stage of the disease. Two EGFP-positive cells are observed fully enveloping individual neurons, highlighted by circles.
Neuronal nuclei were visualized with Hoechst staining. Images were acquired over a 3-hour period at a rate of one frame every 5 minutes.

### Slide 14
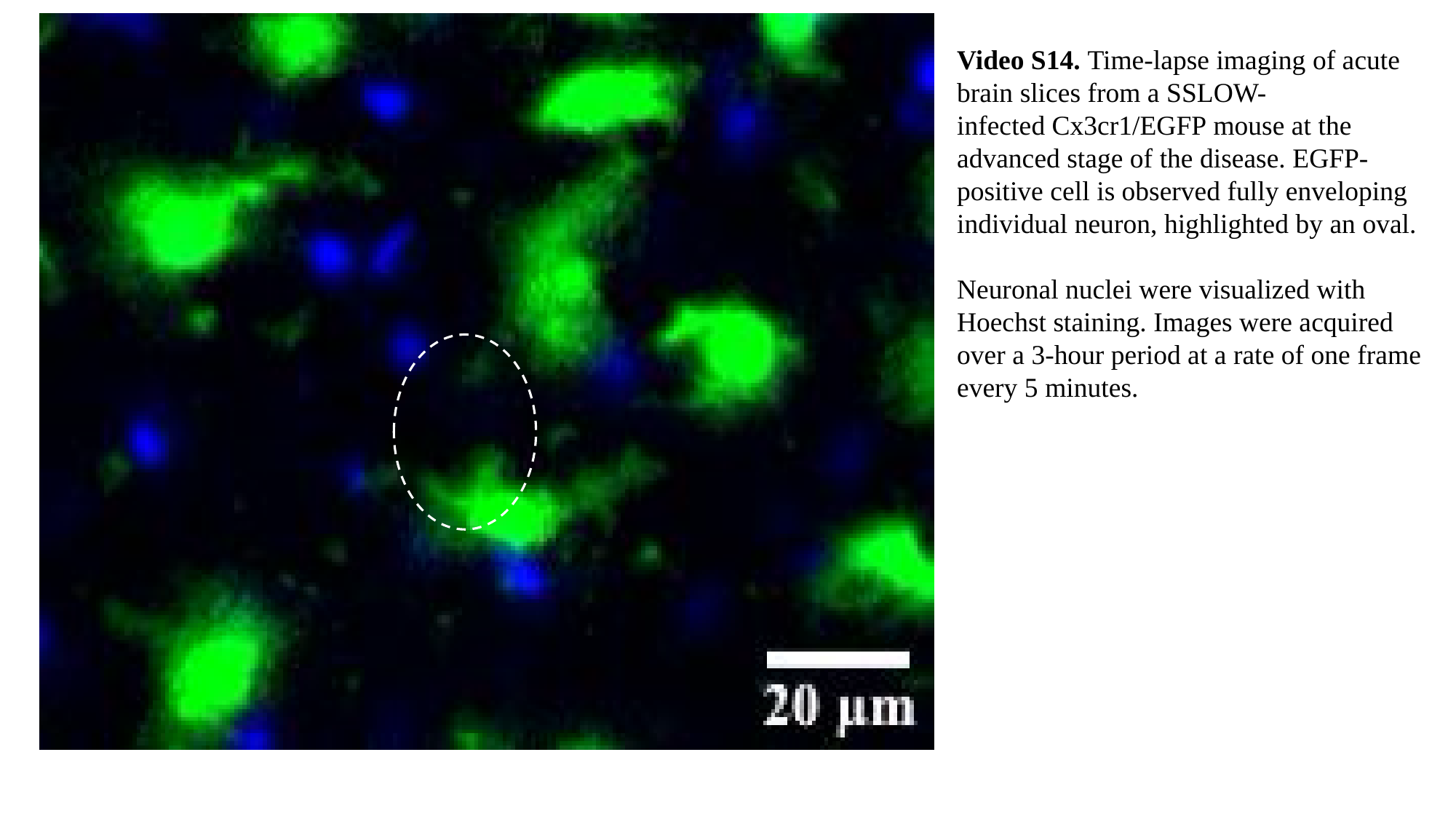

Video S14. Time-lapse imaging of acute brain slices from a SSLOW-infected Cx3cr1/EGFP mouse at the advanced stage of the disease. EGFP-positive cell is observed fully enveloping individual neuron, highlighted by an oval.
Neuronal nuclei were visualized with Hoechst staining. Images were acquired over a 3-hour period at a rate of one frame every 5 minutes.

### Slide 15
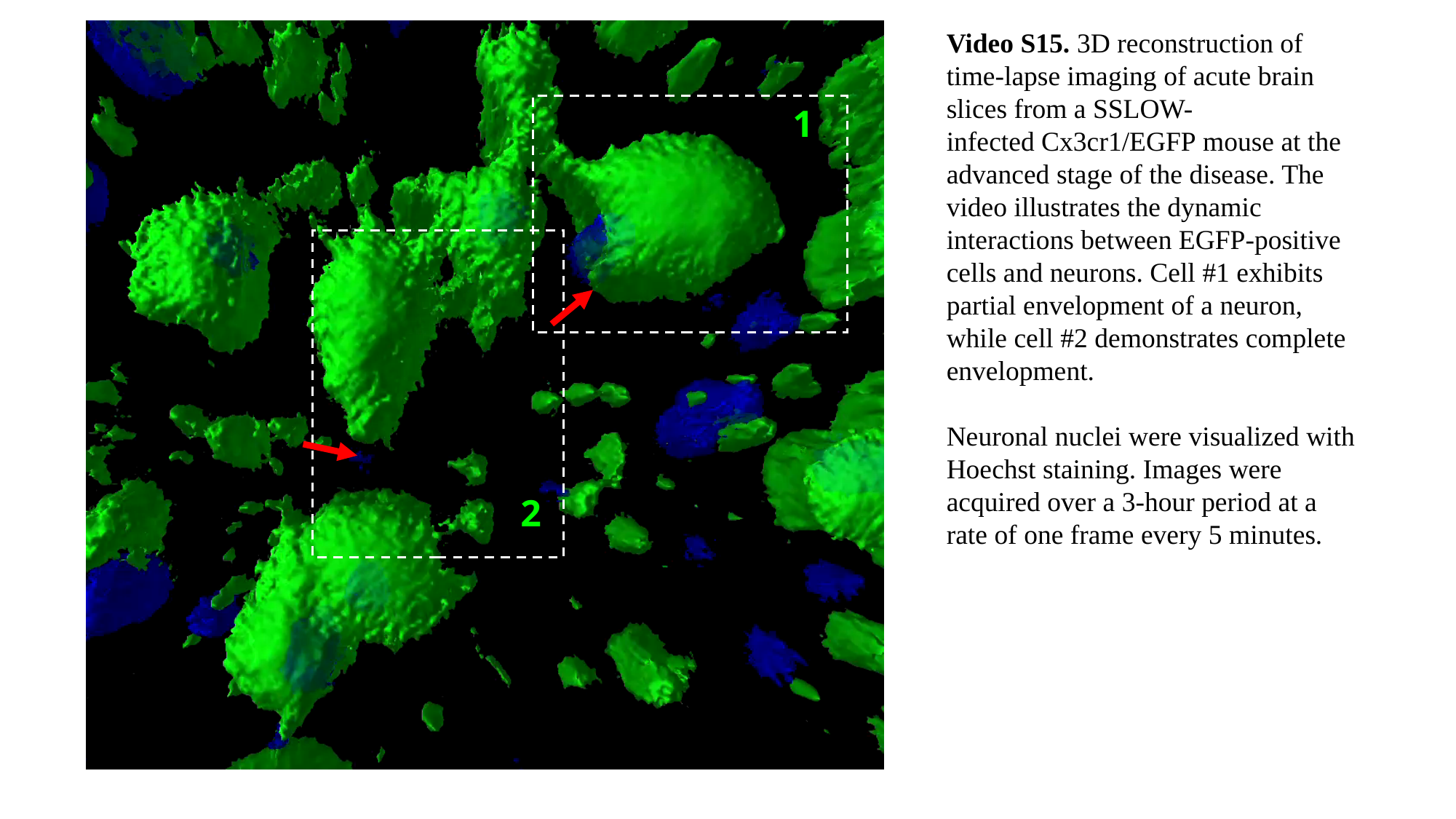

Video S15. 3D reconstruction of time-lapse imaging of acute brain slices from a SSLOW-infected Cx3cr1/EGFP mouse at the advanced stage of the disease. The video illustrates the dynamic interactions between EGFP-positive cells and neurons. Cell #1 exhibits partial envelopment of a neuron, while cell #2 demonstrates complete envelopment.
Neuronal nuclei were visualized with Hoechst staining. Images were acquired over a 3-hour period at a rate of one frame every 5 minutes.
1
2

### Slide 16
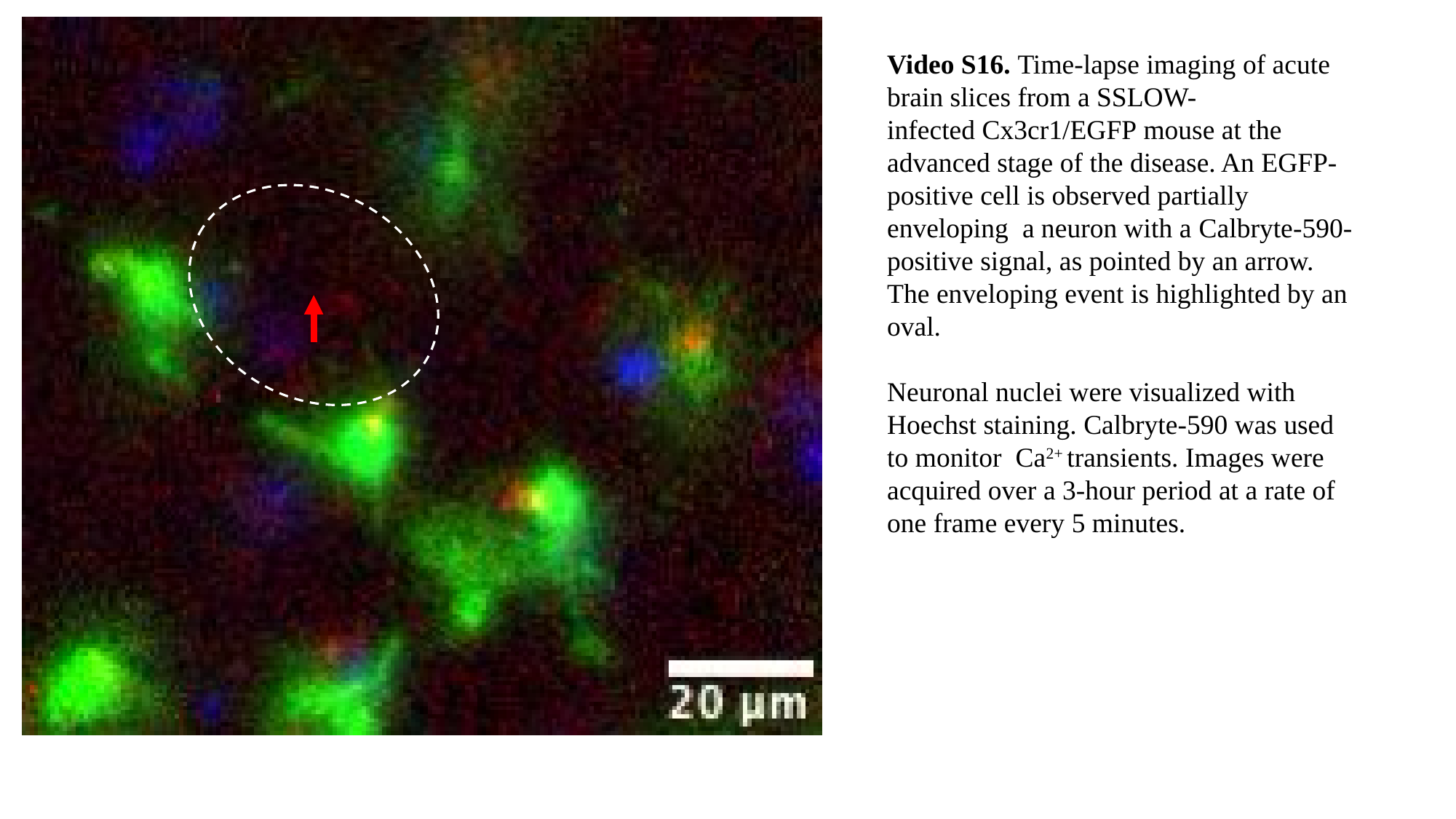

Video S16. Time-lapse imaging of acute brain slices from a SSLOW-infected Cx3cr1/EGFP mouse at the advanced stage of the disease. An EGFP-positive cell is observed partially enveloping a neuron with a Calbryte-590-positive signal, as pointed by an arrow. The enveloping event is highlighted by an oval.
Neuronal nuclei were visualized with Hoechst staining. Calbryte-590 was used to monitor Ca^2+^ transients. Images were acquired over a 3-hour period at a rate of one frame every 5 minutes.

### Slide 17
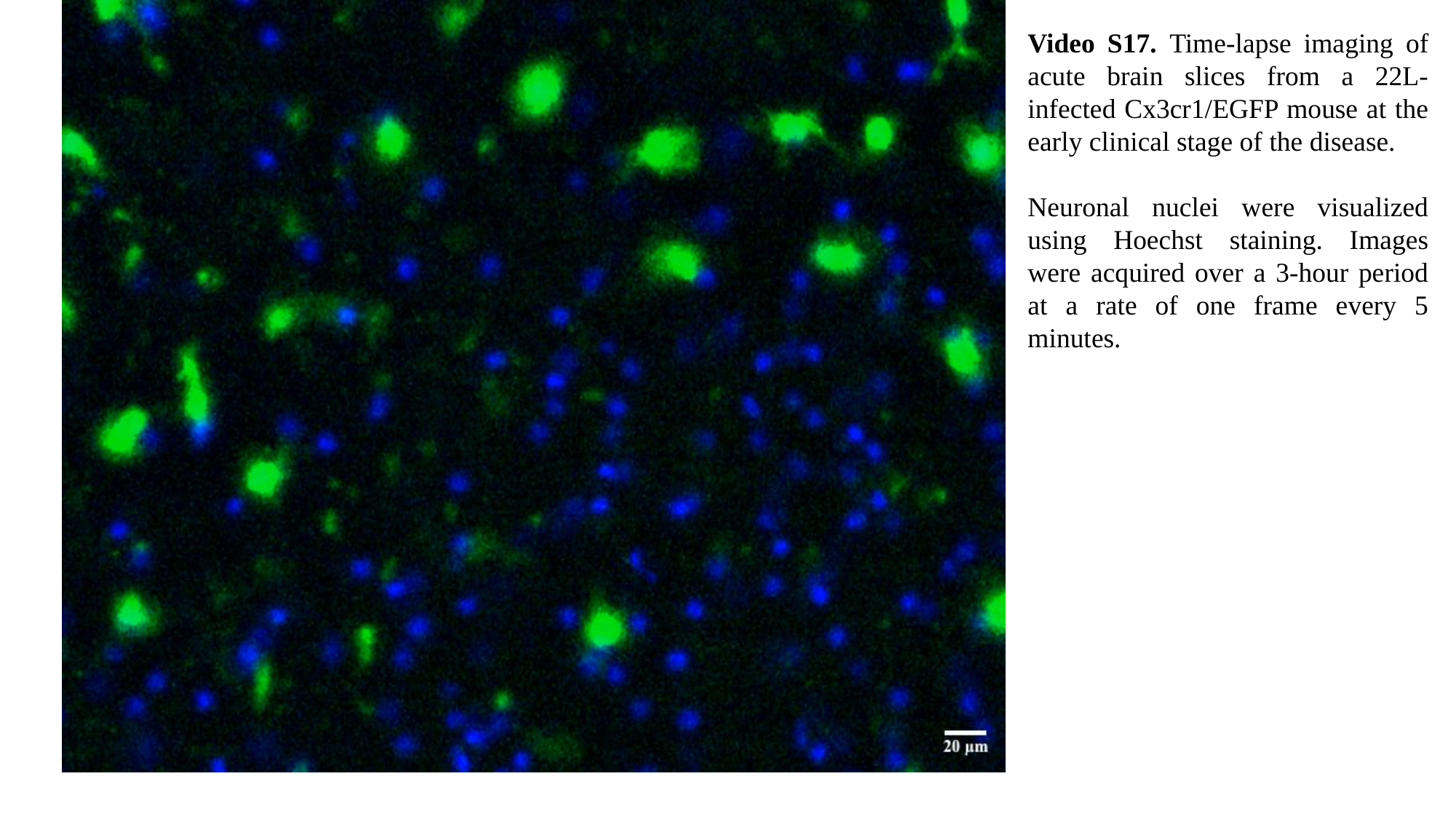

Video S17. Time-lapse imaging of acute brain slices from a 22L-infected Cx3cr1/EGFP mouse at the early clinical stage of the disease.
Neuronal nuclei were visualized using Hoechst staining. Images were acquired over a 3-hour period at a rate of one frame every 5 minutes.

### Slide 18
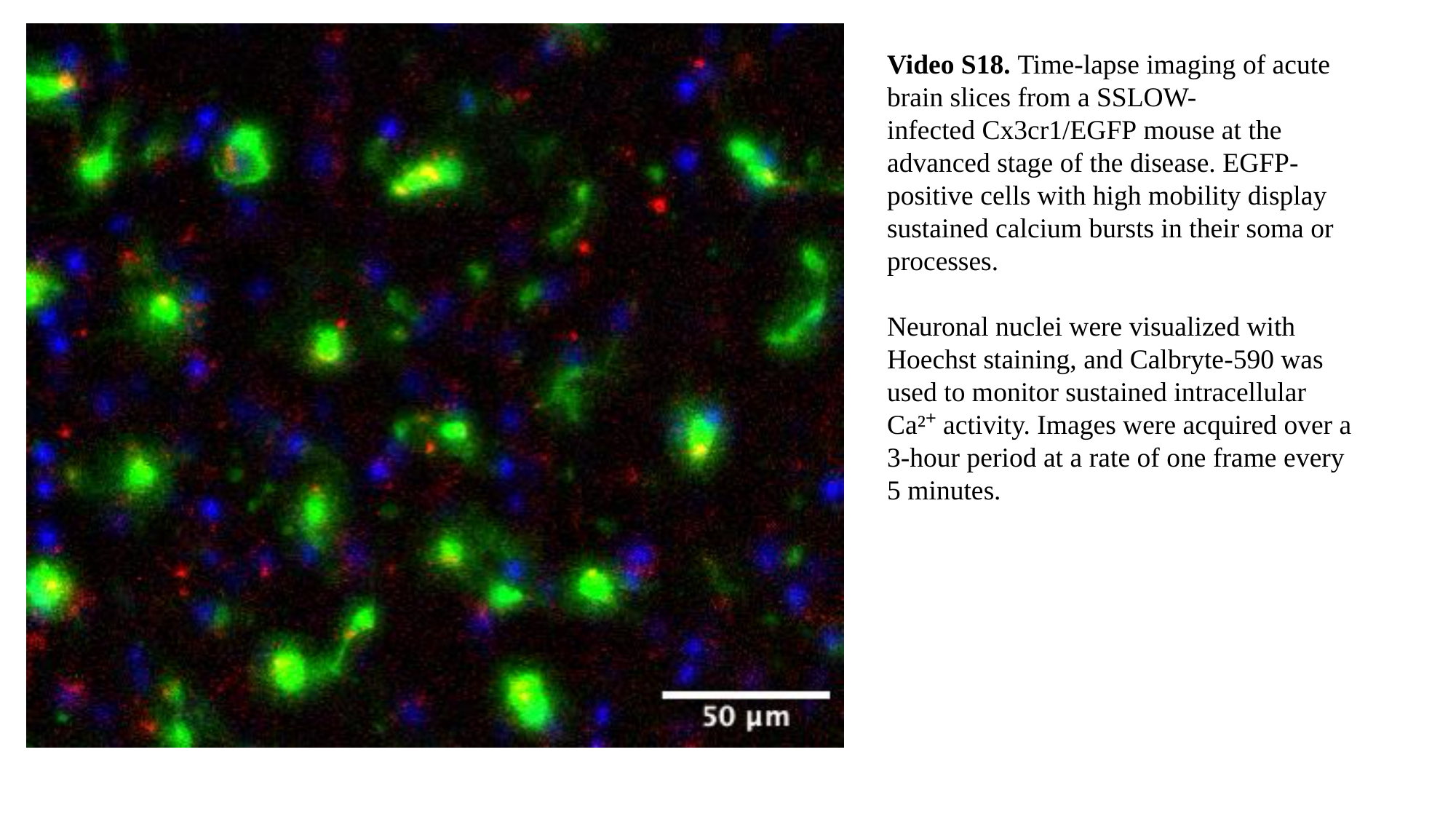

Video S18. Time-lapse imaging of acute brain slices from a SSLOW-infected Cx3cr1/EGFP mouse at the advanced stage of the disease. EGFP-positive cells with high mobility display sustained calcium bursts in their soma or processes.
Neuronal nuclei were visualized with Hoechst staining, and Calbryte-590 was used to monitor sustained intracellular Ca²⁺ activity. Images were acquired over a 3-hour period at a rate of one frame every 5 minutes.

### Slide 19
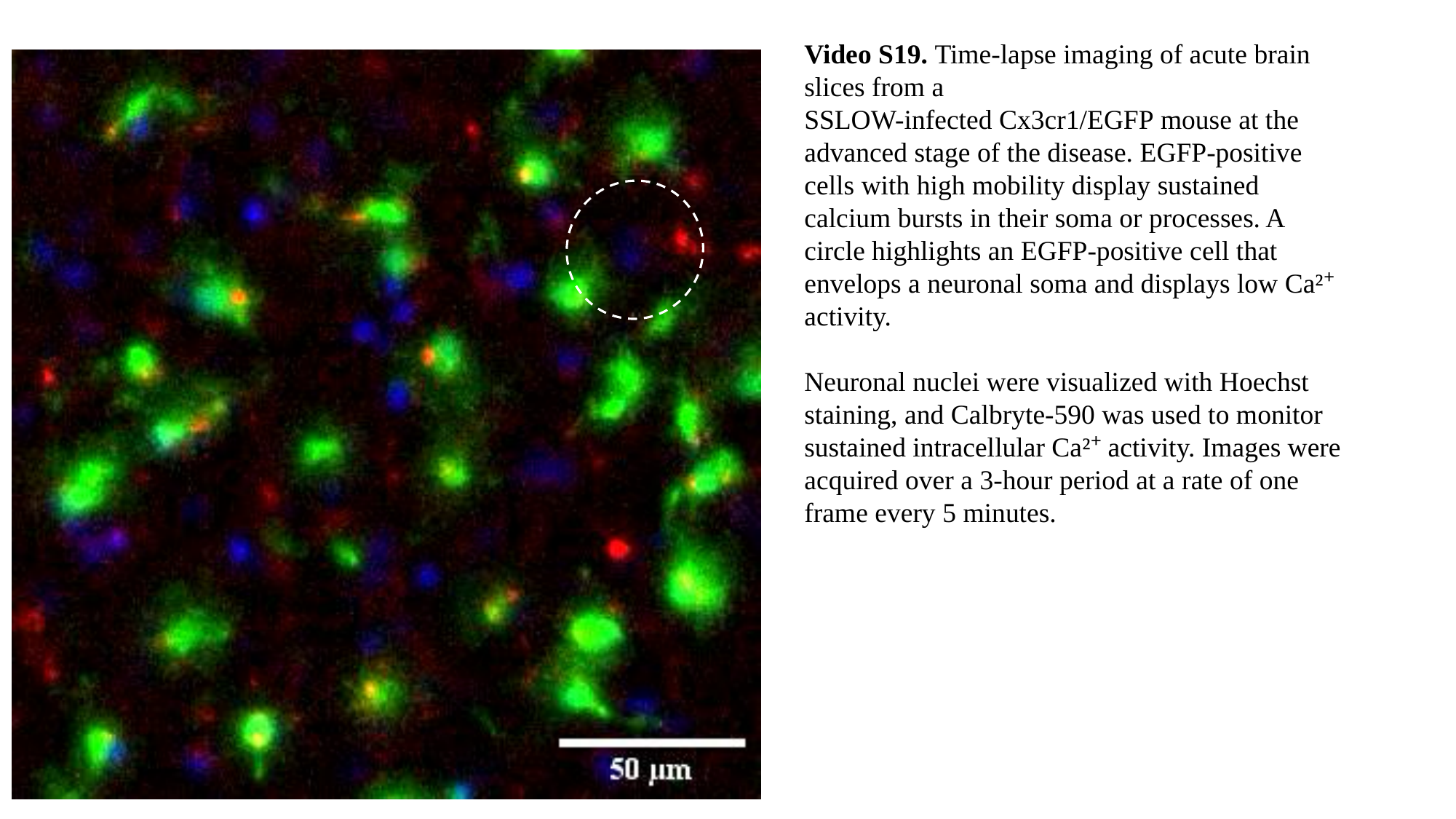

Video S19. Time-lapse imaging of acute brain slices from a SSLOW-infected Cx3cr1/EGFP mouse at the advanced stage of the disease. EGFP-positive cells with high mobility display sustained calcium bursts in their soma or processes. A circle highlights an EGFP-positive cell that envelops a neuronal soma and displays low Ca²⁺ activity.
Neuronal nuclei were visualized with Hoechst staining, and Calbryte-590 was used to monitor sustained intracellular Ca²⁺ activity. Images were acquired over a 3-hour period at a rate of one frame every 5 minutes.

### Slide 20
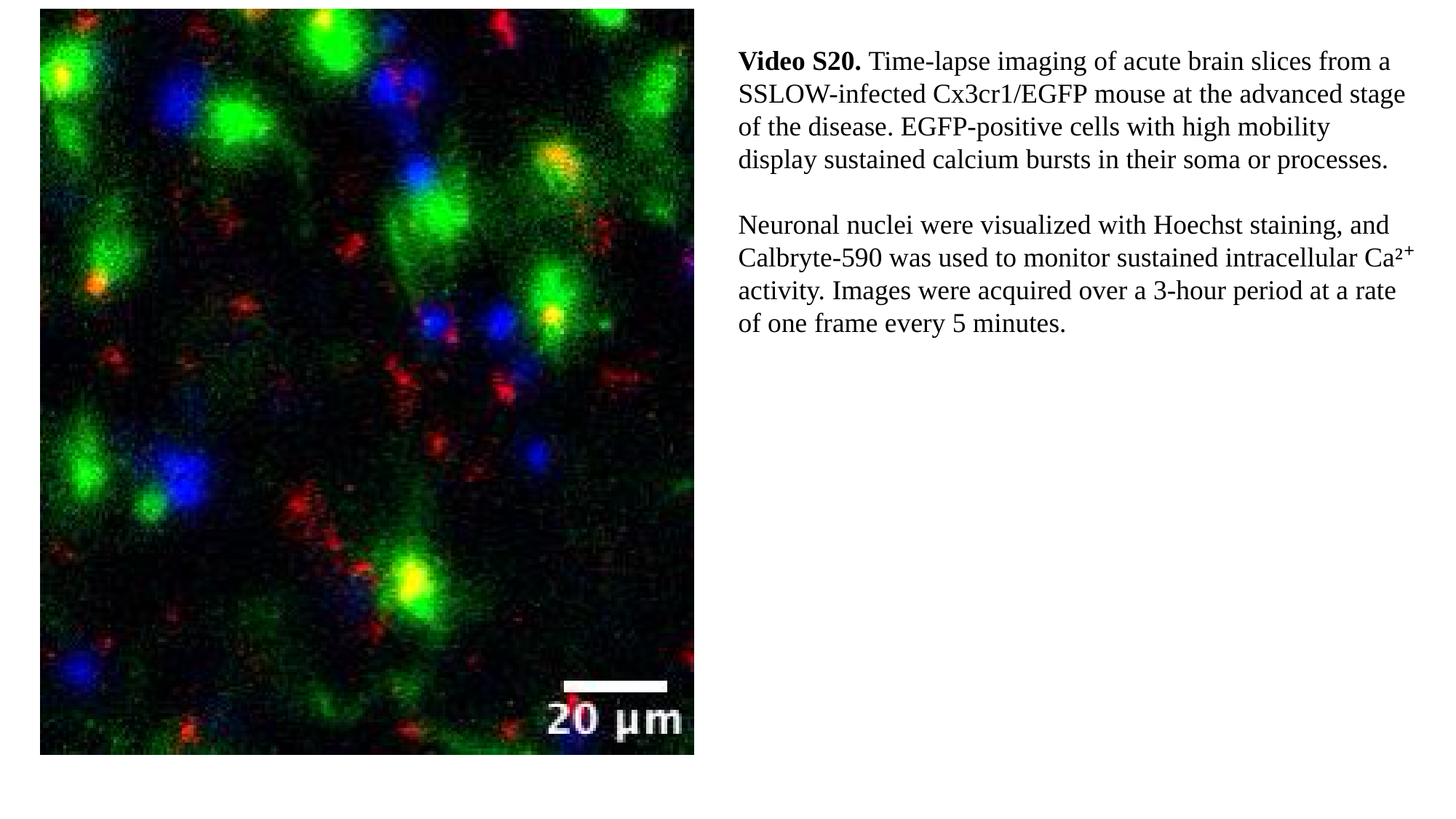

Video S20. Time-lapse imaging of acute brain slices from a SSLOW-infected Cx3cr1/EGFP mouse at the advanced stage of the disease. EGFP-positive cells with high mobility display sustained calcium bursts in their soma or processes.
Neuronal nuclei were visualized with Hoechst staining, and Calbryte-590 was used to monitor sustained intracellular Ca²⁺ activity. Images were acquired over a 3-hour period at a rate of one frame every 5 minutes.

### Slide 21
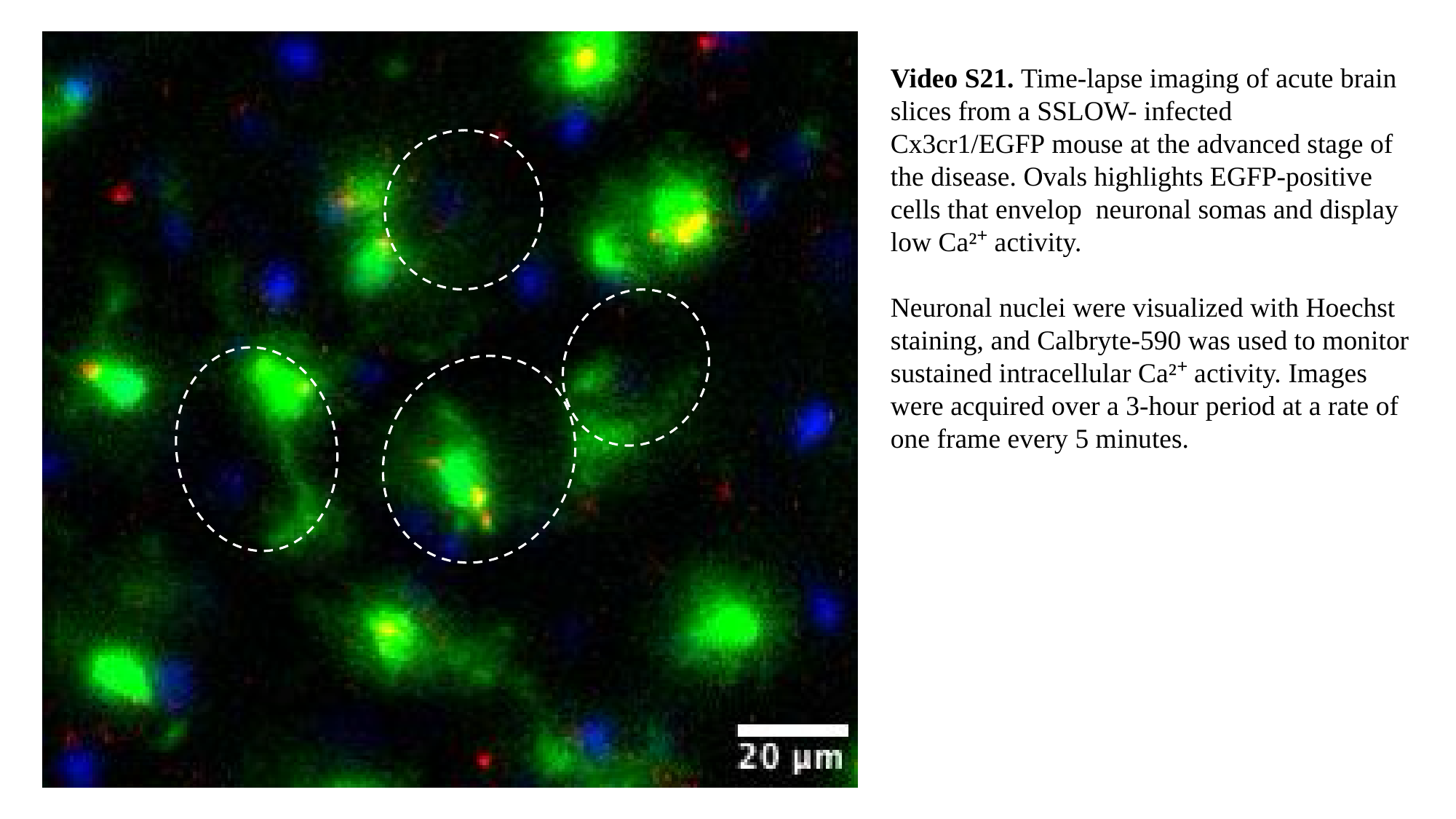

Video S21. Time-lapse imaging of acute brain slices from a SSLOW- infected  Cx3cr1/EGFP mouse at the advanced stage of the disease. Ovals highlights EGFP-positive cells that envelop neuronal somas and display low Ca²⁺ activity.
Neuronal nuclei were visualized with Hoechst staining, and Calbryte-590 was used to monitor sustained intracellular Ca²⁺ activity. Images were acquired over a 3-hour period at a rate of one frame every 5 minutes.

### Slide 22
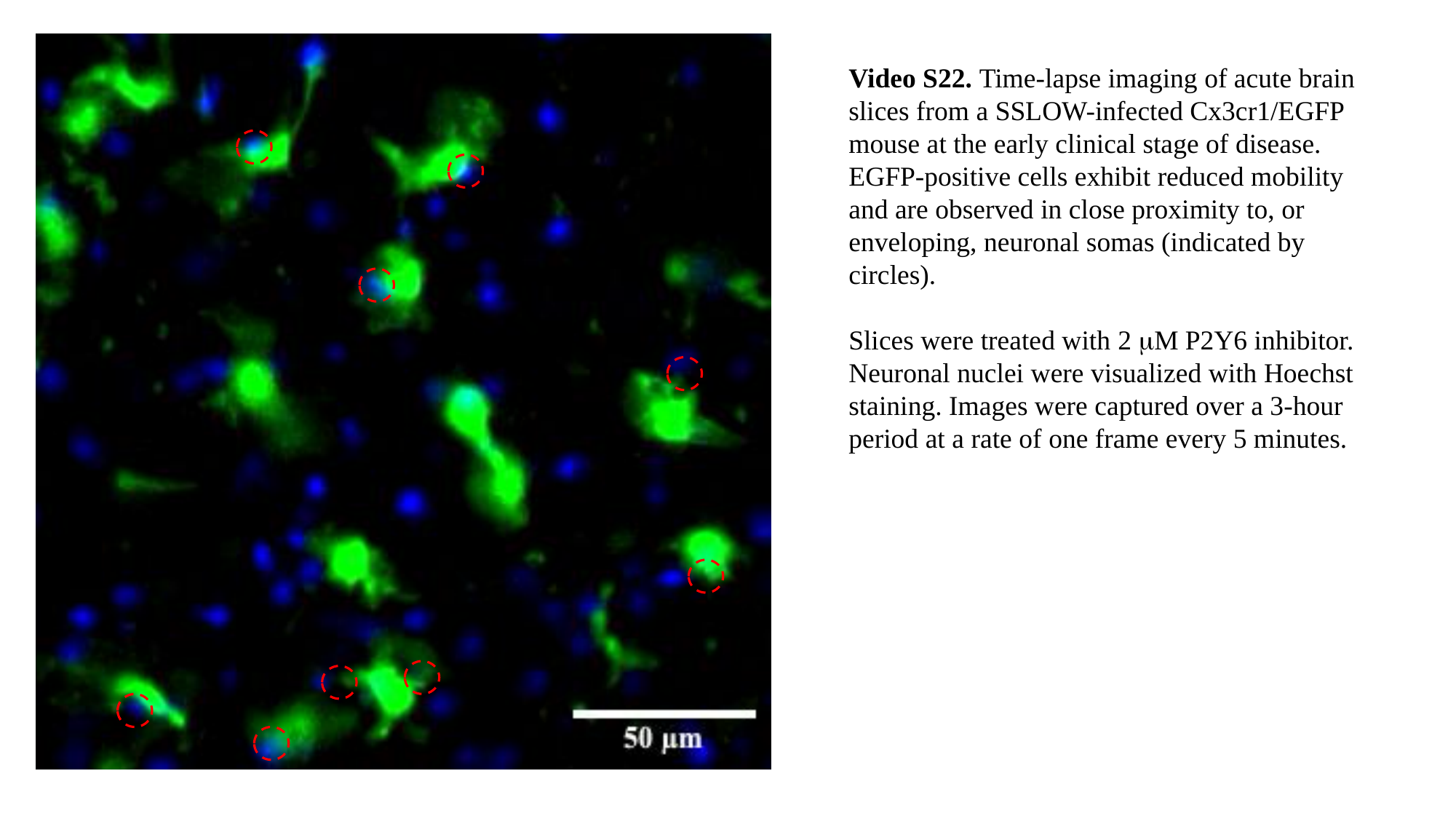

Video S22. Time-lapse imaging of acute brain slices from a SSLOW-infected Cx3cr1/EGFP mouse at the early clinical stage of disease. EGFP-positive cells exhibit reduced mobility and are observed in close proximity to, or enveloping, neuronal somas (indicated by circles).
Slices were treated with 2 μM P2Y6 inhibitor.
Neuronal nuclei were visualized with Hoechst staining. Images were captured over a 3-hour period at a rate of one frame every 5 minutes.

### Slide 23
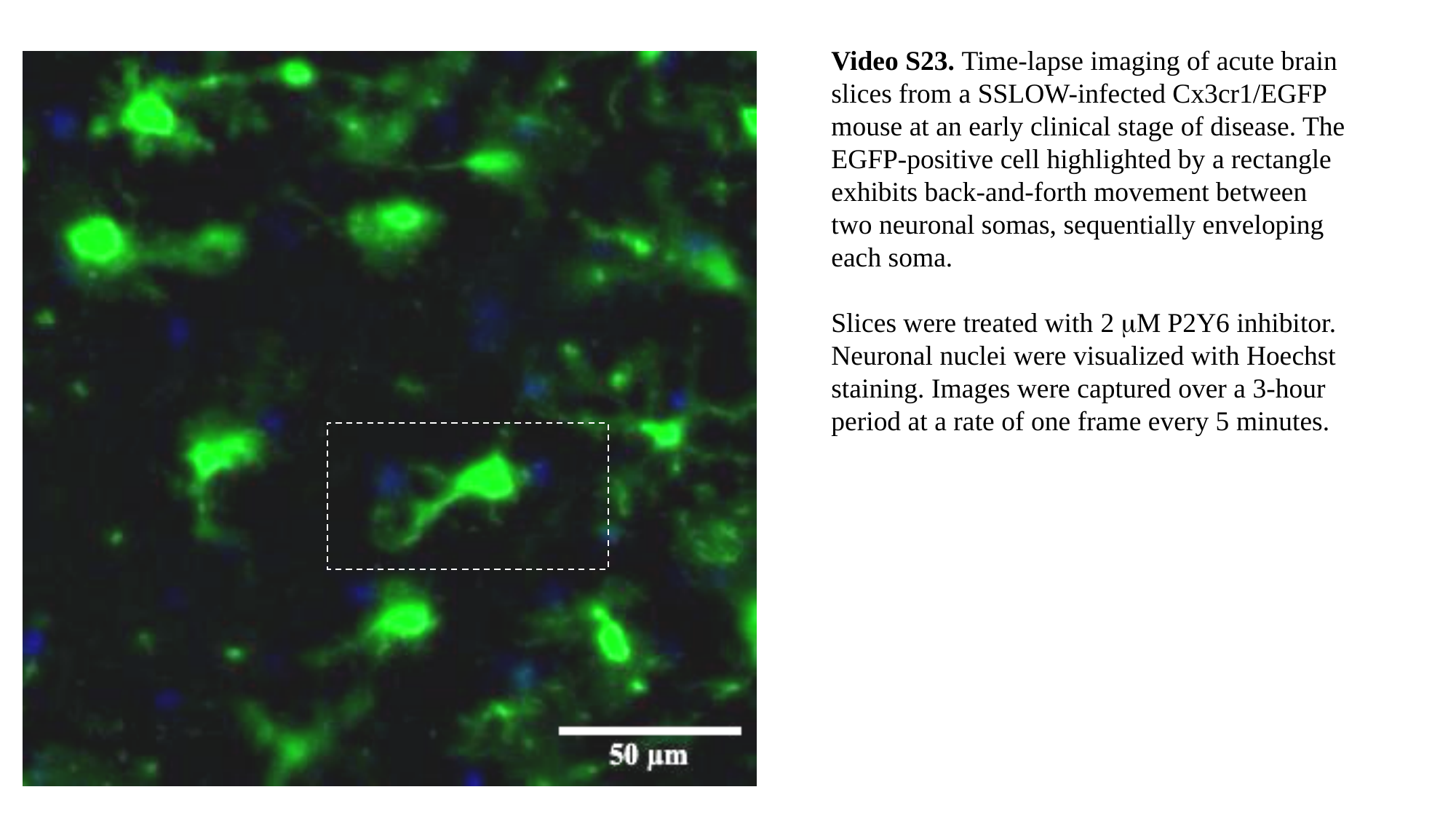

Video S23. Time-lapse imaging of acute brain slices from a SSLOW-infected Cx3cr1/EGFP mouse at an early clinical stage of disease. The EGFP-positive cell highlighted by a rectangle exhibits back-and-forth movement between two neuronal somas, sequentially enveloping each soma.
Slices were treated with 2 μM P2Y6 inhibitor.
Neuronal nuclei were visualized with Hoechst staining. Images were captured over a 3-hour period at a rate of one frame every 5 minutes.
